# Perturbed DNA methylation by sustained overexpression of Gadd45b induces chromatin disorganization, DNA strand breaks and dopaminergic neuron death in mice

**DOI:** 10.1101/2020.06.23.158014

**Authors:** Camille Ravel-Godreuil, Olivia Massiani-Beaudoin, Philippe Mailly, Alain Prochiantz, Rajiv L. Joshi, Julia Fuchs

**Affiliations:** Center for Interdisciplinary Research in Biology (CIRB), Collège de France, CNRS, INSERM, Université PSL, Paris, France; Orion Imaging Facility, Center for Interdisciplinary Research in Biology (CIRB), Collège de France, CNRS, INSERM, Labex Memolife, Université PSL, Paris, France

## Abstract

Heterochromatin disorganization is a key hallmark of aging and DNA methylation state is currently the main molecular predictor of chronological age. The most frequent neurodegenerative diseases like Parkinson disease and Alzheimer’s disease are age-related but how the aging process and chromatin alterations are linked to neurodegeneration is unknown. Here, we investigated the consequences of viral overexpression of *Gadd45b*, a multifactorial protein involved in active DNA demethylation, in the midbrain of wild-type mice. *Gadd45b* overexpression induces global and stable changes in DNA methylation, particularly on gene bodies of genes related to neuronal functions. DNA methylation changes were accompanied by perturbed H3K9me3-marked heterochromatin and increased DNA damage. Prolonged *Gadd45b* expression resulted in dopaminergic neuron degeneration accompanied by altered expression of candidate genes related to heterochromatin maintenance, DNA methylation or Parkinson disease. *Gadd45b* overexpression rendered midbrain dopaminergic neurons more vulnerable to acute oxidative stress. Heterochromatin disorganization and DNA demethylation resulted in derepression of mostly young LINE-1 transposable elements, a potential source of DNA damage, prior to *Gadd45b*-induced neurodegeneration. Our data implicate that alterations in DNA methylation and heterochromatin organization, LINE-1 derepression and DNA damage can represent important contributors in the pathogenic mechanisms of dopaminergic neuron degeneration with potential implications for Parkinson disease.

## Introduction

Epigenetic marks separate chromatin into actively transcribed euchromatin and repressive heterochromatin domains, and participate in the spatial organization of the genome into highly structured 3D domains ^1^. These epigenetic signatures include DNA methylation, various post-translational modifications of histones, and attraction forces between different types of genomic repeat elements, including transposable elements (TEs) ^1^. Changes in chromatin structure regulate DNA accessibility to transcription factors and the physical proximity of enhancers to promoters, thereby regulating gene expression.

It is well established that DNA methylation regulates various important cellular processes during development and cell differentiation. In recent years, however, perturbations of chromatin organization have been linked to the aging process and global changes in DNA methylation are currently the main molecular predictor of chronological age (reviewed in ^2^). Aging-induced epigenetic remodeling of chromatin can impact on genomic stability ^3,4^, and vice versa ^2^. Thus chromatin states and genomic stability are important, interdependent factors associated with the aging process ^5^. One emerging culprit related to both processes is the un-silencing of transposable elements (TEs) with age. Around half of the human genome is comprised of TEs. The evolutionary most successful TEs in mammals are long interspersed nuclear element-1 (LINE-1 or L1). Mostly fossilized and a few remaining active LINE-1 sequences (around 100 in humans and 3000 in mice) represent around 17% of the human ^6^ and 21% of the mouse genome ^7^. Young and full-length LINE-1 elements are autonomous retrotransposons, expanding in the genome through a “copy and paste” retrotransposition mechanism and encoding the two necessary proteins, namely ORF1p and ORF2p, required for their mobilization. ORF1p is an RNA binding protein with strong “cis” preference ^8–10^ and ORF2p encodes an endonuclease and a reverse transcriptase ^11,12^. Several repressive cellular mechanisms, including DNA methylation and heterochromatinization, limit their expression ^13^. When these fail with age, TEs can become derepressed ^14^. An increased activity of LINE-1 is associated with genomic instability through the induction of DNA damage ^15–18^.

How aging and neurodegeneration are linked at the molecular level remains widely unknown. This question is, however, of high relevance as age is the primary risk factor for the most common neurodegenerative diseases (NDs) including Parkinson disease (PD) and Alzheimer disease (AD) ^19^. Some of the cellular processes defined as the hallmarks of aging ^20^ overlap with pathways shown to be dysfunctional in NDs. This is the case for impaired proteostasis, mitochondrial dysfunction, deregulated nutrient sensing, increased oxidative stress and neuroinflammation ^21, 22^. Whether two other important nominators of aging, namely perturbations in the chromatin organization and genomic instability, are associated with neuronal aging and neurodegeneration has not been unequivocally proven yet.

A cardinal feature of PD is the degeneration of midbrain dopaminergic (mDA) neurons in the substantia nigra *pars compacta* (SNpc). These neurons project to the striatum and their loss leads to a striatal deficiency in the neurotransmitter dopamine, inducing the typical motor symptoms of PD. Decreased global DNA methylation with age in the SNpc has been observed ^23^ and DNA methylation changes, mostly on specific genetic risk loci, have been linked to several NDs ^24^ including PD ^25^. Alterations in histone modifications have also been observed in PD ^26^. However, the possible contribution of age-related epigenetic alterations to the pathogenesis of PD and the onset of neurodegeneration has not been demonstrated yet.

Here, we investigated how SNpc mDA neurons react to perturbations of chromatin organization. For this purpose, we overexpressed *Gadd45b* in the SNpc of wild-type mice using an adeno-associated virus (AAV) vector. GADD45B is a multifunctional protein which coordinates in the nucleus an active DNA demethylation pathway involving cytidine deaminases and DNA glycosylases ^27^ in association with the base excision repair (BER) pathway ^28^. We chose GADD45B because it is a known DNA demethylase in postmitotic neurons ^27, 29^ and it is highly inducible in conditions of oxidative stress in the SNpc ^30^. We show that overexpression of *Gadd45b* in the SNpc of wild-type mice leads to widespread perturbations of DNA methylation, heterochromatin disorganization, increased vulnerability of mDA neurons to oxidative stress, activation of LINE-1 elements, DNA strand breaks and neuronal death. Our data reinforces the hypothesis that aging-induced global chromatin disorganization initiates neurodegeneration, possibly via the derepression of LINE-1 elements.

## Results

### Gadd45b overexpression in the SNpc of wild-type mice leads to early and stable DNA methylation perturbations in gene bodies of genes related to neuronal functions

Wild-type mice littermates (6 weeks old) were injected unilaterally in the SNpc using *AAV8-mCherry* or AAV8-*mGadd45b*. The animals were sacrificed after 14 or 90 days *post-injectionem* (p.i.) and either perfused or dissected as schematized in **Figure 1A**. The AAV8-mCherry control virus efficiently infected tyrosine hydroxylase positive (TH+) neurons of the SNpc as shown by mCherry expression at 14d p.i. (***Fig. 1B, upper panel***). Due to the lack of a good antibody against GADD45B, expression of *Gadd45b* was verified at 14d p.i. by *in situ* hybridization (***Fig1. B, lower panel***) and RT-qPCR after manual micro-dissection of the SNpc at 14d and 90d p.i. (***Fig. 1C***). *Gadd45b* transcripts on the injected ipsilateral side were increased up to 79-fold on average (11.02±4.83; 869.1±283; n=6) at 14d p.i. and up to 178-fold (7.89±1.52; 1402±683; n=4) at 90d p.i. compared to the endogenous transcript levels of the non-injected contralateral side.

**Figure 1.**
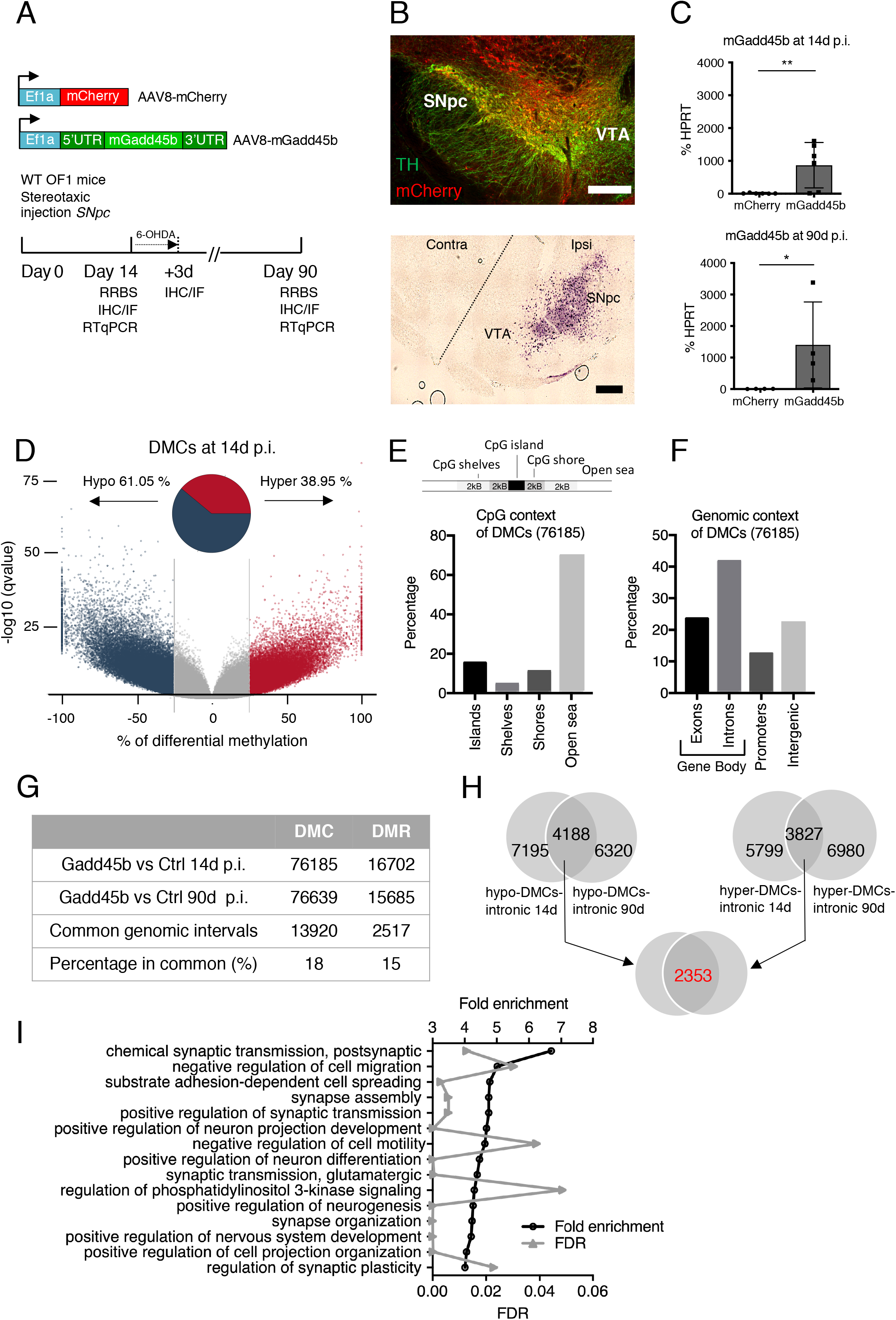
Gadd45b overexpression in the SNpc of wildtype mice leads to perturbed DNA methylation. **A**: Overview of the viral injection protocol into the SNpc of wild-type mice. Wild-type OF1 mice were injected unilaterally in the SNpc with AAV8 virus expressing either mCherry or mouse Gadd45b with its regulatory sequences under the control of the Ef1a promoter. Animals were then sacrificed 14 or 90 days later to perform RRBS, immunostainings or RT-qPCRs. **B**: The AAV8-mCherry virus diffuses within the SNpc and infects mDA neurons. **Upper panel**: TH and mCherry immunostaining of the SNpc of AAV8-mCherry injected mice showing the diffusion of the virus in the SNpc area while sparing the VTA. Scale bar represents 300 μm. **Lower pane**l: ISH of Gadd45b expression. Gadd45b expression in the SNpc of AAV8-mGadd45b injected mice at 14d p.i. shows that Gadd45b is overexpressed only in the ipsilateral SNpc and not in the ipsilateral VTA, nor on the contralateral side. **C:** Exogenous Gadd45b RNA is expressed 14d after the injection of AAV8-mGadd45b. Transcript expression analysis by RT-qPCR following manual microdissection of the SNpc of the injected side (**upper panel**) show a 79-fold increase in mGadd45B transcript level (mCherry= 11.02±4.83; mGadd45B= 869.10+282.80; mean + SEM) at 14d p.i. (left); n=6; and a 178 fold increase (7.89+1.52; 1402+683) at 90d p.i. (right); n=4; *p<0,05; error bars represent SEM. **D-I**: RRBS analysis of differentially methylated CpGs (DMCs) in the SNpc region after injection of AAV8-mGadd45b or AAV8-control. **D**: Volcano plot of differentially methylated CpGs shows widespread perturbations in DNA methylation upon Gadd45b overexpression 14d p.i.. 76 185 significantly differentially methylated CpGs were detected with a q-value smaller than or equal to 0.01 and at least 25% difference. The Volcano plot shows the number of regions with changed patterns of methylation between Gadd45b and control samples significantly higher or lower than the 25% difference cut-off and considering a q-value threshold of 0.01. The difference in methylation is reflected in the x-axis while the y-axis represents the significance of the difference. Regions that are highly differentially methylated are further to the left and right sides of the plot, while highly significant changes appear higher on the plot. Values on the x- and y-axes are percent methylation differences and negative log10 of the corrected p-values, respectively. The pie chart shows the percentage of hyper- and hypomethylated regions. **E and F**: DMCs are located majorly in open sea regions and in introns of genes. DMCs were annotated in relation to the distance to a CpG island (E), as well as based on the genomic regions they are associated with (**F**). Their distributions are plotted in a bar chart. **G**: Comparison of total number, common genetic intervals (minimum overlap of 1 bp) and percentage in common for DMCs and DMRs 14d and 90d p.i.. **H**: The “Gadd45b-DMC-regulon”. Venn diagrams showing the overlap of genes containing at least one intronic hypomethylated DMC at 14 and 90d (4188 genes, left upper Venn diagram), the overlap of genes containing at least one hypermethylated DMC at 14d and 90d (3827 genes, right upper Venn diagram) and the overlap of both groups (2353 genes, lower Venn diagram) representing the “Gadd45b-DMC-regulon”. Note that the “Gadd45b-DMR-regulon” comprising 447 genes is summarized in the **Suppl. Fig.1E**. **I**: GO-analysis of the “Gadd45b-DMC-regulon” reveals neuron-related gene categories enriched after Gadd45b overexpression 14d p.i.. Gene ontology analysis (PANTHER version 15.0) and the PANTHER overrepresentation test with the GO-Slim annotation data set ‘biological process’ identified significantly overrepresented GO categories. The first fifteen significantly overrepresented GO categories with the highest fold enrichment are displayed with the fold enrichment on the left y-axis (black points) and the FDR value on the right y-axis (grey points).

Having verified the efficient overexpression of *Gadd45b* in the SNpc, we asked whether this perturbs DNA methylation patterns in the SNpc. To this end, we injected wild-type mice with *AAV8-mGadd45b* (n=6) or *AAV8-mCherry* (n=6) and simultaneously extracted DNA and RNA from manually micro-dissected SNpc biopsies at 14d p.i. (***Scheme Fig. 1A***). The DNA of two mice per condition, selected for high expression by RT-qPCR of *Gadd45b* or *mCherry,* respectively, was then subjected to reduced representation bisulfite sequencing (RRBS) for DNA methylation analysis. Bioinformatic analysis of RRBS data detected 809117 CpGs that were common between both conditions. Of those, 76185 individual CpGs were differentially methylated (DMCs) with a q-value smaller than, or equal to, 0.01 and at least 25% difference. Using a window and step size of 1000 bp, differentially methylated regions (DMRs) were defined and 16702 regions passed the defined significance threshold. A majority of DMCs and DMRs were hypomethylated throughout time, located in open sea regions (defined as regions outside CpG islands, CpG shores or CpG shelves) and localized in gene bodies, particularly introns. The Volcano plot in ***Figure 1D*** illustrates the DMC methylation pattern at 14d p.i.. Upon *Gadd45b* overexpression, 46508 DMCs (61.05 %) were hypomethylated and 29677 DMCs (38.95%) were hypermethylated. The percentage of hypo- and hypermethylated regions per chromosome was similar (***Suppl. Fig. 1G***). We then interrogated DMCs in relation to their distance to a CpG island and found 69.73% to be overlapping with open sea regions (***Fig. 1E**),* defined as regions outside of a known CpG island, CpG shore (2000 bp flanking the CpG island) or CpG shelves (2000 bp flanking the CpG shores). Analysis of the genomic context revealed that 41.36% of DMCs were located in introns (***Fig. 1F***), followed by 22.3% in the intergenic space, 23.6% in exons and 12.5% in promoter regions. This analysis indicates early and widespread methylation changes primarily in gene bodies upon *Gadd45b* overexpression.

In order to understand the long-term changes in methylation patterns induced by *Gadd45b* overexpression, we extracted DNA from mice 90 days after injection of AAV8-*mGadd45b* or AAV8-*mCherry* control. Analysis revealed a similar distribution of hypo- and hypermethylated DMRs at 14d p.i. (***Suppl. Fig. 1A**)* and at 90d p.i. (***Suppl. Fig. 1C***). The genomic context of DMCs and DMRs, both at 14d p.i. (***Suppl. Fig. 1B**)* and at 90d p.i. (***Suppl. Fig. 1D**)* was very similar as well, the majority of differential methylation concerning open sea regions and gene bodies, particularly introns (summarized in ***Table 1***). The overall numbers of DMRs and DMCs with AAV8-*mGadd45b* also resembled that at 14d p.i., but common DMRs or DMCs examination revealed an overlap of only 15% of DMRs (2517) and 18% of DMCs (13920) (***Fig. 1G***). While this indicates that the specific regions with methylation changes induced by *Gadd45b* overexpression are not stable over time, the general location in open sea regions and in introns of genes of DMRs and DMCs is maintained as a specific and stable footprint of *Gadd45b* overexpression, which we termed “*Gadd45b*-regulon”. Furthermore, there was an important overlap of more than half of the genes containing intronic hypomethylated or hypermethylated DMCs at 14d and at 90d (***Fig. 1H***). The extent of overlap was similar for intronic DMRs (hypoDMRs at 14d: 3937 genes; hypoDMRs at 90d: 3467 genes, overlap (1712 common genes); ***Suppl. Fig. 1E***). Of those common genes, 2353 genes contained at least one intronic hypo- and one intronic hypermethylated DMC (***Fig. 1H**)* and 447 genes at least one intronic hypo- and one intronic hypermethylated DMR at both 14d and 90d p.i. (***Suppl. Fig. 1E***). Thus, *Gadd45b* overexpression induces stable changes in methylation patterns in gene bodies, particularly introns.

**Table 1.** Summary of the differential methylation analysis comparing Gadd45b overexpression to mCherry control in the SNpc of wild-type mice.

| Differential methylation analysis<br>Gadd45b/ mCherry | 14d p.i. |  | 90d p.i. |  |
| --- | --- | --- | --- | --- |
|  | DMCs | DMRs | DMCs | DMRs |
| Total number | 76185 | 16702 | 76639 | 15685 |
| Hypomethylated (in %) | **61,05** | **64,57** | **54,50** | **56,35** |
| **Distance to CpG island (in %)** |  |  |  |  |
| Islands | 15,34 | 1,50 | 16,03 | 1,30 |
| Shelves | 4,23 | 6,30 | 4,18 | 6,60 |
| Shores | 10,71 | 7,60 | 10,73 | 7,60 |
| Open sea | **69,73** | **84,60** | **69,06** | **84,60** |
| **Association with genomic regions (in %)** |  |  |  |  |
| Exons | **23,60** | **19,22** | **24,00** | **19,06** |
| Introns | **41,60** | **45,39** | **41,40** | **45,03** |
| Promoters | 12,50 | 8,57 | 12,80 | 9,03 |
| Intergenic | 22,30 | 26,83 | 21,90 | 26,88 |

The results of the RRBS analysis prompted us to identify functional categories associated with this GADD45B-induced methylation footprint corresponding to the 2353 genes stably containing DMCs over time (***Fig. 1H***). We used Gene ontology analysis (PANTHER version 15.0; http://pantherdb.org) and the PANTHER overrepresentation test with the GO-Slim annotation data set ‘biological process’. 145 GO categories were significantly overrepresented with a fold change >2 and an FDR <0.05. The first fifteen significantly overrepresented GO categories are displayed in ***Figure 1I***. Of those, 10 GO categories are explicitly related to neuronal functions with two prevailing categories, namely synaptic function and organization, and neurodevelopment and neurogenesis. This is very similar to what we found for DMRs (***Suppl. Fig. 1F***) and suggests that *Gadd45b* is involved in the specific regulation of gene body methylation of neuron-related genes.

### Gadd45b overexpression leads to heterochromatin disorganization in mDA neurons

Members of GADD45 protein family have been described to promote heterochromatin relaxation ^31^. We therefore examined whether *Gadd45b* overexpression, in addition to global methylation changes, would also alter chromatin organization, in particular the organization of heterochromatin. To do so, we stained mDA neurons for histone H3 lysine 9 trimethylation (H3K9me3), a repressive heterochromatin mark. Immunostaining for H3K9me3 shows a perinucleolar pattern composed, on average, of 3 or 4 foci (3.64±0.12) in TH+ neurons in the SNpc of AAV8-*mCherry* injected mice (***Fig. 2A***). This pattern becomes disorganized in AAV8-*mGadd45b* injected mice at 14d p.i.. Semi-automated quantification of H3K9me3 staining specifically in TH+ neurons identified a 1.13-fold increase in the number of H3K9me3 foci (4.10±0.12) scattered across the nucleus, an increase by 1.24-fold of the average H3K9me3 foci volume (4.78±0.15; 5.91±0.22 μm^3^), a reduction by 1.21 fold in the intensity of the diffuse nucleoplasmic H3K9me3 staining (2.37×10^7^±557961; 1.96×10^7^±441292), but no difference in foci intensity (1.32×10^6^±54236; 1.41×10^6^±64942) (***Fig. 2A***, quantification in ***Fig. 2B-E***). This shift in heterochromatin organization is still detectable at 90d p.i. (***Fig. 2F-J***). These results show that a de-structuration of heterochromatin is already detectable 14d and stable up to 90d after the injection of AAV8-*mGadd45b* indicating an early and stable perturbation of global heterochromatin organization upon *Gadd45b* overexpression.

**Figure 2.**
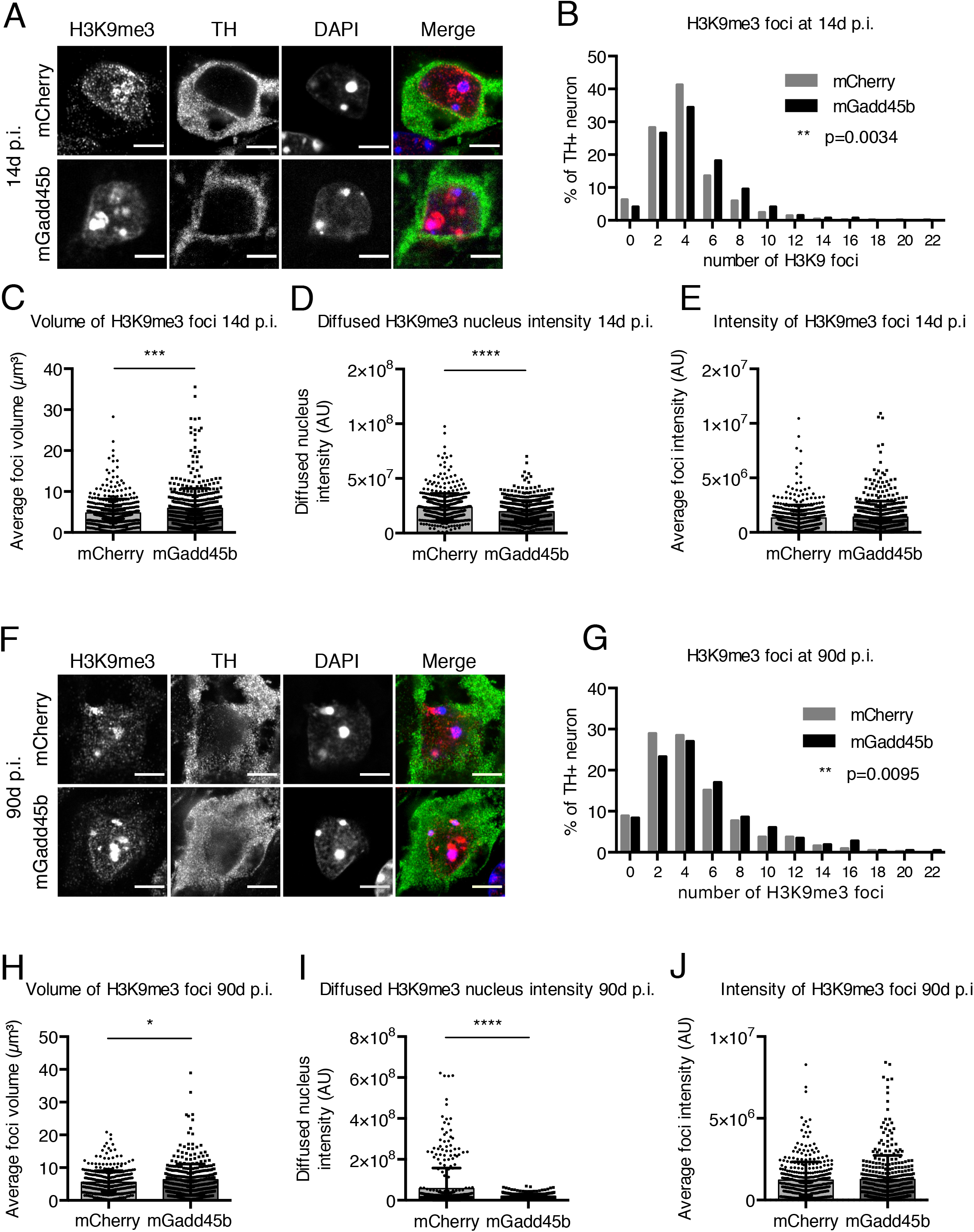
Gadd45b overexpression leads to heterochromatin destructuration in mDA neurons. **A-E:** H3K9me3 heterochromatin staining is perturbed as early as 14d after the injection of AAV8-mGadd4b. TH+ neurons in the SNpc of AAV8-mGadd45b injected mice (**A**) display at 14d p.i. a 1.13-fold increase in the number (3.64±0.12; 4.10±0.12, data represented as a frequency distribution histogramm in **B**) and a 1.24-fold increase in the volume (4.78±0.15; 5.91±0.22 μm^3^, **C**) of H3K9me3 foci. The diffuse nucleoplasmic staining intensity (2.37×10^7^±557961; 1.96×10^7^±441292) is decreased by 1.21-fold (**D**) while the foci intensity (1.32×10^6^±54236; 1.41×10^6^±64942) remains unchanged (**E**). **F-J:** H3K9me3 heterochromatin staining remains perturbed up until 90d after the injection of AAV8-mGadd45b. A similar pattern as in **A-E** was observed at 90d p.i. with a 1.18-fold increase in the number (4.07+0.17; 4.82+0.20, data represented as a frequency distribution histogramm in **G**) and a 1.15-fold increase in the volume (5.51+0.18; 6.35+0.24 μm^3^, **H**) of H3K9me3 foci shown in (**F**). The diffuse nucleoplasmic staining intensity (5.66×10^7^+5×10^6^; 1.97×10^7^+573406) is decreased by 2.87-fold while the foci intensity (1.22×10^6^+56943; 1.26×10^6^+74064) remains unchanged (**I**; **J**). Scale bar in **A** and **F** represents 5 μm. ** p<0.01; *** p<0.001; **** p<0.0001; n=3 mice; Between 510 and 534 neurons were quantified per condition at 14d p.i. and 428 neurons were quantified per condition at 90d p.i.. Error bars represent SEM.

We next examined whether *Gadd45b* overexpression leads to perturbations in the pattern of the DNA methylation marker MeCP2. Immunostaining of sections from mice injected with AAV8-*mGadd45b* or *AAV8-mCherry* did not show any difference in the number of MeCP2 foci in TH+ neurons (***Suppl. Fig. 2A-D***), the intensity of the diffuse nucleoplasmic MeCP2 staining nor the volume and intensity of MeCP2 foci, neither at 14d nor at 90d p.i. (***Suppl. Fig. 2E,G-J***). There was a slight increase in foci intensity at 14d p.i. with AAV8-*mGadd45b (**Suppl. Fig. 2F***). This analysis indicates that DNA methylation perturbation as detected by RRBS does not imply a global change in the organization of MeCP2 distribution.

### Gadd45b overexpression leads to the loss of TH+ neurons

To examine the effect of *Gadd45b* overexpression on the survival of mDA neurons in the SNpc, we quantified the number of TH+ neurons in mice injected with *AAV8-mCherry* or *AAV8-mGadd45b* at 14d and 90d p.i. (***Fig. 3A***). There was no difference between TH+ cell numbers comparing AAV8-*mCherry*- and AAV8-*mGadd45b*-injected (ipsilateral) sides to the non-injected side (contralateral) at 14d p.i. (***Fig. 3B**: AAV8-mCherry* ipsi/contra: 1.04±0.02; AAV8-*mGadd45b* ipsi/contra: 0.97±0.04). However, 90d after injection of *AAV8-mGadd45b,* this ratio shifted to an average of 0.82±0.04, indicating a specific loss of 18% of TH+ neurons on the AAV8-m*Gadd45b* injected side compared to the contralateral side (***Fig. 3C***). The TH+ neurons ratio between the ipsi- and the contralateral side remained unchanged in AAV8-*mCherry* injected mice (***Fig. 3C**:* ipsi/contra: 0.97±0.02). These results show that *Gadd45b* overexpression in the SNpc can trigger degeneration of mDA neurons in the long-term. In addition, the absence of a significant TH+ cell loss at 14d p.i. suggests that alterations in the distribution of methylation at CpGs and in the organization of chromatin precede neuronal death.

**Figure 3.**
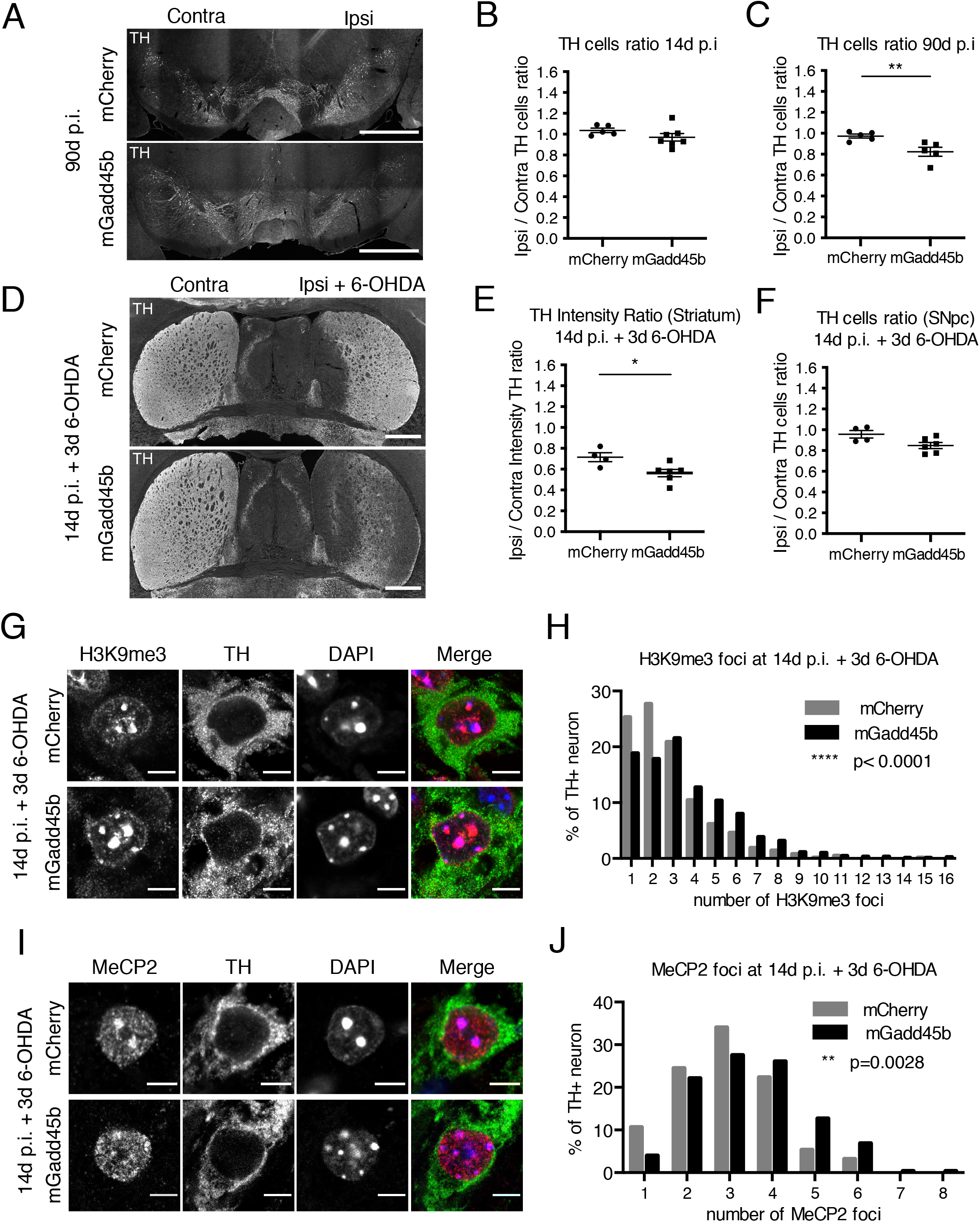
Loss of TH+ neurons and their increased vulnerability upon Gadd45b overexpression. **A-C**: Dopaminergic cell loss in the SNpc 90d following the injection of AAV8-mGadd45b. The ratio of the number of TH+ neurons of the injected side on the non-injected side is similar at 14d p.i. between AAV8-mCherry and AAV8-mGadd4b injected mice and close to 1 (1.04+0.02; 0.97+0.04) (**B**). At 90d p.i. however, if the ratio of AAV8-mCherry mice is still close to 1 (0.97±0.02), the ratio of AAV8-mGadd45b mice (0.82±0.04) reflect a 18 % loss of TH+ neurons compared to the non-injected side, shown in (**A**) and quantified in (**C**); scale bar represents 1000 μm; n=5. **D-F**: Dopaminergic neurons are sensitized to 6-OHDA upon Gadd45b overexpression. At 14d p.i., AAV8-mCherry and AAV8-mGadd45b mice were injected with 6-OHDA in the striatum, on the same side as the viral injection and analyzed 3 days after (**Fig. 1A**). TH staining of dopaminergic axons shows that axonal degeneration in the striatum induced by 6-OHDA occurs in AAV8-mCherry mice (0.71±0.04) but is 1.5-fold more pronounced in AAV8-mGadd45b mice (0.56±0.04) shown in (**D**), quantified in (**E**). The number of TH+ neurons is unaffected (0.96+0.04; 0.85+0.03) after the 3 day 6-OHDA treatment (**F**); n= 4-6; Scale bar represents 1000 μm. **G-J**: Gadd45b expression accentuates heterochromatin loss and MeCP2 foci dispersion upon striatal 6-OHDA injection. TH+ neurons in the SNpc of AAV8-mGadd45b mice injected with 6-OHDA at 14d p.i. display 3 days after the 6-OHDA injection an increase by 1.47-fold in the number of H3K9me3 foci (3.28±0.15; 4.82±0.16), shown in (**G**), quantified and represented as a frequency distribution histogramm in (**H)** and by 1.16-fold in the number of MeCP2 foci (2.97±0.12; 3.45±0.08), shown in (**I**), quantified and represented as a frequency distribution histogramm in (**J**). Scale bar represents 5 μm; * p<0,05; ** p<0,01; **** p<0,0001; n=3; Between 632 and 887 neurons were quantified per condition for H3K9me3 foci and between 94 and 276 neurons per condition for MeCP2 foci. Error bars represent SEM.

### Enhanced vulnerability of mDA neurons overexpressing Gadd45b to oxidative stress

To examine whether the heterochromatin de-structuration observed in TH+ neurons at 14d p.i. upon *Gadd45b* overexpression could render these neurons more vulnerable to oxidative stress, mice were first injected with AAV8-*mCherry* or AAV8-*mGadd45b* and 14d later with 6-hydroxy-dopamine (6-OHDA, 2 μl; 0.5 μg/μl) in the ipsilateral striatum. When injected into the striatum, 6-OHDA induces a specific and retrograde death of mDA neurons in the SNpc and is frequently used to model PD in rodents ^32^ including mice ^33^. Three days after the unilateral, striatal injection of 6-OHDA, immunostainings of striatal sections show a 29% loss (mCherry ipsi/contra: 0.71±0.04) of TH intensity in the ipsilateral striatum compared to the contralateral side in mCherry expressing mice (illustrated in ***Fig. 3D*,** quantified in **Fig. 3E**). This decrease in TH staining intensity reaches 44% (mGadd45b ipsi/contra: 0.56±0.04) in AAV8-*mGadd45b* injected mice (illustrated in ***Fig. 3D*,** quantified in **Fig. 3E**), suggesting that *Gadd45b* overexpression increases the axonal degeneration of mDA neurons induced by 6-OHDA. However, this experimental paradigm did not lead to any significant loss of TH cell bodies in the SNpc (***Fig. 3F**),* potentially due to the time of analysis. We analyzed mice 3 days rather than 6 days after 6-OHDA injection, the time point normally used to induce mDA cell death in the SNpc ^32^, to identify early events in TH+ cell bodies. An increased vulnerability to oxidative stress of mDA neurons overexpressing *Gadd45b* was also reflected at the heterochromatin level. The injection of 6-OHDA changed the nuclear localization and increased the number of H3K9me3 positive foci (***Fig. 3G, H**)* compared to AAV8-*mCherry*. Under these conditions, the number of MeCP2 positive foci in mDA neurons expressing AAV8-*mGadd45b* was also increased as compared to *AAV8-mCherry (**Fig. 3I, J***). This presumably reflects more significant DNA methylation changes upon *Gadd45b* overexpression under oxidative stress.

### Chromatin de-structuration is accompanied by increased DNA damage

The heterochromatin loss model of aging stipulates that heterochromatin de-condensation is a driving force of cellular aging ^34^. Loss of proteins involved in heterochromatin maintenance has been shown to lead to increased DNA damage and accelerated aging ^35, 36^. In the context of NDs, recent studies have reported that chromatin relaxation in the brain could also result in increased DNA damage and genome instability ^18, 37–39^. Since *Gadd45b* overexpression led to chromatin changes, we investigated whether it might also induce DNA damage. We therefore performed immunostainings for phosphorylated histone H2AX (γ-H2AX), a marker for DNA strand breaks (***Fig. 4A***). TH+ neurons contained either a single perinucleolar focus or a diffuse nucleoplasmic staining without any foci. After injection of AAV8-*mGadd45b* (14d p.i.), the majority (65.94±2.10 %) of TH+ neurons displayed a diffuse, intense nucleoplasmic staining against only 39.81±2.40 % of TH+ neurons after *AAV8-mCherry* injection, the majority showing one or more prominent perinucleolar γ-H2AX-positive foci (***Fig. 4B, C***). The quantification of the diffuse γ-H2AX staining in the nucleus revealed a more intense staining after *Gadd45b* overexpression, indicating widespread DNA damage.

**Figure 4.**
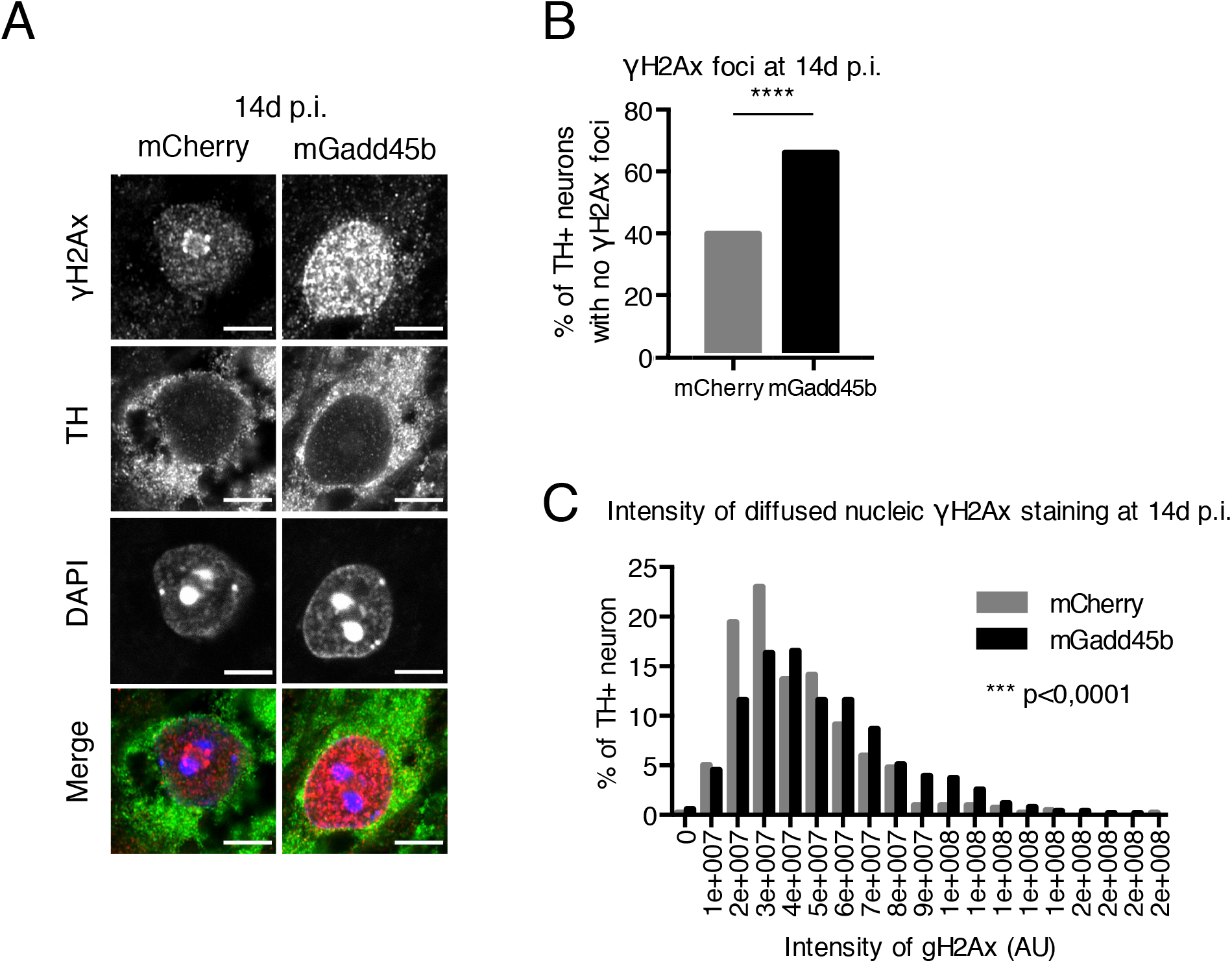
DNA methylation changes and heterochromatin de-structuration upon Gadd45b overexpression are associated with DNA damage. **A-C:** Gadd45b overexpression induced DNA damage in TH+ neurons of the SNpc. The majority (65.94+2.10 %) of TH+ neurons of the SNpc in AAV8-mGadd45b display no single γ-H2AX foci but instead an intense diffuse nucleoplasmic staining, 1.7-times more than in AAV8-mCherry mice (39.81+2.40 %), shown in (**A**), number quantified in (**B**), diffuse nucleoplasmic intensity quantified and represented as a frequency distribution histogram in (**C**). Scale bar represents 5 μm; n=3. Between 417 and 518 neurons were quantified per condition.

### LINE-1 methylation is affected by Gadd45b overexpression

Heterochromatin alterations can unsilence normally repressed TEs, including LINE-1 elements ^14^. LINE-1 are a potential source of DNA damage ^16-18, 40^. Repeat elements are detected by RRBS and the Bismark software used for mapping of the RRBS reads only considers uniquely mapped reads to avoid any bias during the methylation calling. Thus, reads that are mapping with a same mapping score to multiple locations on the reference genome, which will be the case for most reads derived from repetitive elements, will not be considered. We exploited these facts to explore the methylation status of LINE-1 elements in our experimental conditions. Using the RRBS data, we used the LINE-1 annotation included in the software HOMER, which contains its own database of LINE-1 elements for the mouse genome to interrogate the overlap of DMRs and DMCs with LINE-1 elements 14 days after injection of AAV8 viruses. This analysis showed that 3530 DMCs and 1030 DMRs overlapped with an annotated LINE-1 element, which we termed L1-DMCs and L1-DMRs respectively. 794 (22.5%) L1-DMCs (***Fig. 5A**)* and 264 (25.6%) L1-DMRs (***Suppl. Fig. 3A***) were located in intronic regions, and 2734 (77.5%) L1-DMCs (***Fig. 5A***) and 766 L1-DMRs (***Suppl. Fig. 3A**)* in intergenic regions. Of all the L1-DMCs the majority was hypomethylated (1933, 54,8%, ***Fig. 5B***), similarly to L1-DMRs (595, 57,8%).

**Figure 5.**
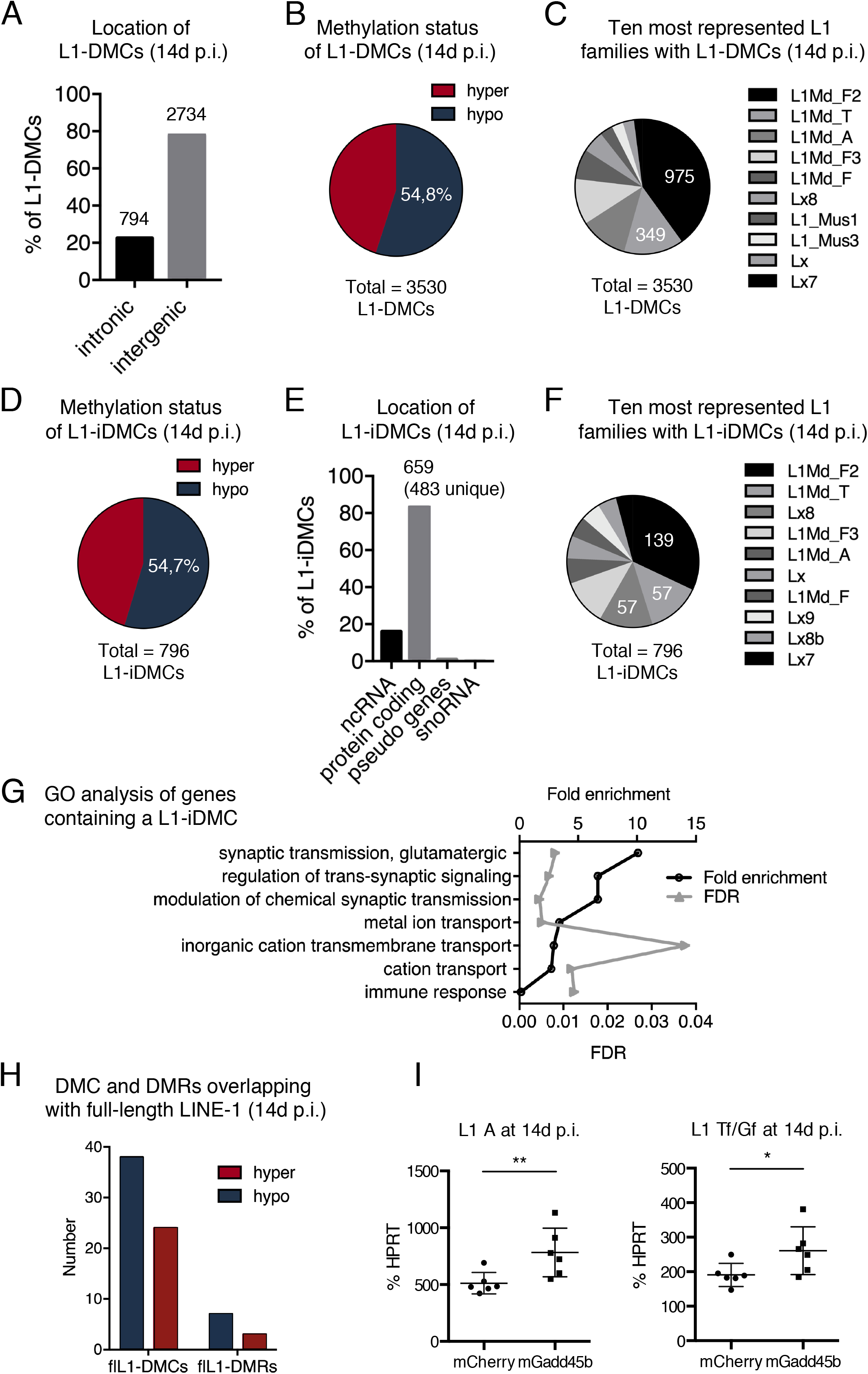
LINE-1 methylation changes and increased LINE-1 expression following Gadd45b overexpression. **A-C:** Analysis of DMCs in LINE-1 sequences (L1-DMCs) 14d p.i. of AAV8-mGadd45b. **A**: Location of L1-DMCs. 22.5% of all L1-DMCs are located in introns. **B**: Of all the L1-DMCs at 14d p.i., most (54.8%) are hypomethylated (blue). **C**: The ten most represented L1 families with at least one L1-DMC. **D-G:** Analysis of DMCs in intronic LINE-1 sequences (L1-iDMCs) 14d p.i. of AAV8-mGadd45b. Most L1-iDMCs (54.7%) are hypomethylated (**D**), located in protein-coding genes (in 483 genes, **E**) and related to neuronal functions (**G**). **F**: The ten most represented L1 families with L1-iDMCs. **H**: Analysis of the methylation status of full-length LINE-1 (flL1) elements containing a DMC or a DMR. Most DMCs and DMRs in full-length L1 elements are hypomethylated. **I:** LINE-1 transcripts of the young L1Md-T and L1Md-A families are increased after injection of AAV8-mGadd45b compared to AAV8-mCherry 14d p.i.. RT-qPCR of RNA extracted from manually micro-dissected SNpc of the injected sides shows a 1.5-fold increase in L1 A transcripts level (**left**) (512.40+38.48; 782.60+87.44) and 1.4-fold increase in L1 Tf/Gf transcripts level (**right**) (190.80+13.63; 261.20+28.28) in AAV8-mGadd45B mice at 14d p.i. compared to AAV8-mCherry mice; *p<0,05; ** p<0,01; **** p<0,0001; n=6. Error bars represent SEM.

To determine which LINE-1 families containing at least one L1-DMC upon *Gadd45b* overexpression were overrepresented, we counted the number of families and ordered them by frequency. ***Figure 5C*** shows the ten most represented LINE-1 families with a L1-DMC. Interestingly, of 94 different families present, three LINE-1 family members, namely L1-Md-F2 (975 L1-DMCs), L1Md-T (349 L1-DMCs) and L1Md-A (280 L1-DMCs), were the three most frequently represented. L1Md-T, L1Md-A and L1-Md-F2 elements are young LINE-1 elements ^41^, which contain full-length, retrotransposition-competent LINE-1 copies. Of LINE-1 associated intronic DMCs (L1-iDMCs, 794) at 14d p.i., the majority (54.7%) was hypomethylated (434 hypomethylated vs 360 hypermethylated, ***Fig. 5D***). L1-iDMCs were mostly located in protein-coding genes (82.8%, ***Fig. 5E***) and the GO analysis of biological processes of 483 genes, containing at least one L1-iDMC, identified seven enriched categories. Among those categories, three were neuron-related (***Fig. 5G***). The most frequent LINE-1 families in intronic L1-iDMCs were, again, the young L1Md-F2 family (139 of 794) and L1Md-T (57 L1-iDMCs) followed by the old Lx8 family (57 L1-iDMCs) (***Fig. 5F***). Notably, other young LINE-1 families were also overlapping with iDMCs (L1Md-A: 27 L1-iDMCs, L1Md-Gf: 6 L1-iDMCs, not shown) (***Fig. 5F***). Similarly to L1-iDMCs, the majority of LINE-1 associated DMRs in introns (L1-iDMRs) was hypomethylated (63,3 %, 167 L1-iDMRs; ***Suppl. Fig. 3A***) and belonged to the L1Md-F2 family (39 of 264) (***Suppl. Fig. 3B***). Hypo- and hypermethylated iDMRs were also found in LINE-1 elements of the active L1Md-T (3 and 2, respectively) and L1Md-A families (3 and 3 elements, respectively). This prompted to examine a data-base annotating full-length LINE-1 elements (L1Basev2 ^42^) to see if intronic and intergenic DMRs and DMCs could coincide with possibly active LINE-1. Ten DMRs and 62 DMCs overlapped with a full-length LINE-1. More than half of them (7 out of 10 DMRs and 38 out of 62 DMCs, ***Fig. 5H, Suppl. Fig. 3C***), were hypomethylated. This data indicates a widespread change in the methylation status of LINE-1 elements upon *Gadd45b* overexpression (14d p.i.). The most differentially methylated LINE-1 elements are hypomethylated and belong to young L1 families (L1Md-F2, L1Md-T, L1Md-A) suggesting a possible expression of these individual LINE-1 loci. Interestingly, L1-iDMCs are frequently located in introns of genes related to neuronal functions (***Fig. 5G***).

### LINE-1 transcripts are increased upon overexpression of Gadd45b

Having established a change in the methylation pattern of LINE-1 elements after injection of AAV8-*mGadd45b*, some of which were full-length, we evaluated the expression of the youngest L1 families in mice, namely L1Md-A and L1Md-Tf/Gf. The analysis by RT-qPCR with specific primers located in the 5’UTR of the L1Md-A (***Fig. 5I, left panel**)* and L1Md-Gf/Tf families (***Fig. 5I, right panel**)* showed a 1.5-fold increase in LINE-1 transcripts in the SNpc after 14d of *Gadd45b* overexpression.

### Gadd45b overexpression is accompanied by expression changes in genes with DMCs

Expression levels of candidate genes with intronic DMCs at either 14d or 90d p.i. were then analyzed by RT-qPCR. We selected 13 genes based on their known function either in chromatin remodeling (*Satb1, Setdb1, Wapl*), DNA methylation (*Tet2, Tet3, Dnmt3a*), PD relevance (*Lrrk2, Park2*), synaptic remodeling (*Sorcs2*), DNA damage (*Xpa*), or in aging and senescence (*Cdkn2a-p19, Cdkn2d, Sirt1*). None of the candidate genes showed a change in expression at 14d p.i. However, at 90d p.i., the expression of *Satb1, Setdb1, Dnmt3a, Tet3* and *Park2* decreased significantly in the ipsilateral SNpc injected with *AAV8-mGadd45b* compared to AAV8-*mCherry*. (**Fig. 6A**). Of note, 4 out of these 5 are genes belonging to the *Gadd45b*-DMC-regulon (**Fig. 1H**). Using RNA-seq data of laser-microdissected SNpc from wild-type mice (GEO *GSE72321*;^30^ and unpublished), we compared the expression levels of the selected candidate genes to those of dopaminergic neuron-specific markers such as *TH*, dopamine transporter *Slc6a3* or the homeogene, *Engrailed* 1 (*En1*)^43^. We found high levels of expression of *Satb1* compared to the other selected genes, indicating that *Satb1* is strongly expressed in the SNpc (**Suppl. Fig. 4**). Exonic and intronic DMCs in downregulated genes comprised hypo-as well as hypermethylated CpGs. Disruption of the normal gene body methylation state of several genes of the GADD45B-DMC-regulon might thus provoke their dysregulation over time but is not associated with a loss of a particular methylation pattern.

**Figure 6:**
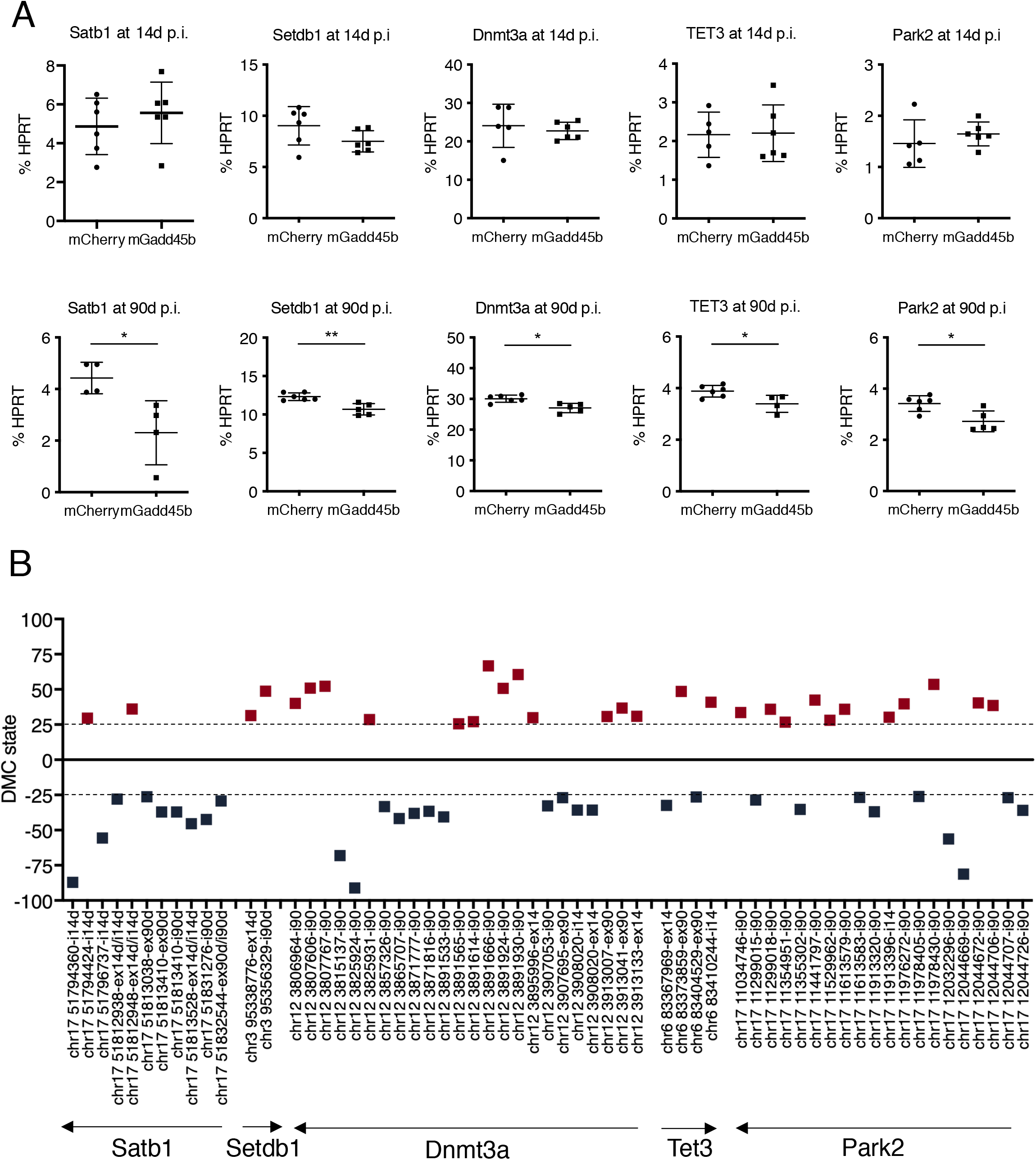
Candidate genes with DMCs upon Gadd45b overexpression show altered expression in the SNpc at 90d p.i. **A:** The expression of several gene candidates containing DMCs is dysregulated. Setdb1, Park2 Dnmt3a, Tet3 and Satb1 transcripts are downregulated 90d p.i. of AAV8-mGadd45b. The mRNA analysis in SNpc of the injected side by RT-qPCR shows no significant difference in Satb1 (4.87+0.59; 5.57+0.65), Setdb1 (9.03+0.76; 7.51+0.42), Dnmt3a (24.07+2.52; 22.76+0.92), Tet3 (2.16+0.26; 2.20+0.65) and Park2 (1.46+0.21; 1.65+0.09) transcripts at 14d p.i. (4.87+0.59; 5.57+0.65); n=5-6. At 90d p.i., there is a decrease in transcript levels of 1.9-fold for Satb 1 (4.43+0.30; 2.31+0.62), of 1.15-fold for Setdb1 (12.32+0.20; 10.68+0.34) of 1.11-fold for Dnmt3a (30.06+0.47; 27.05+0.67), of 1.14 fold for Tet3 (3.89+0.09; 3.40+0.17) of 1.25-fold for Park2 (3.42+0.12; 2.73+0.18), in AAV8-mGadd45b mice compared to AAV8-mCherry mice; * p<0,05; n=4. Error bars represent SEM. **B:** Genes significantly deregulated upon Gadd45b overexpression were analyzed with regard to methylation differences throughout their gene bodies (exonic and intronic DMCs) at 14d and 90d p.i.. Results are displayed for Satb1, Setdb1, Dnmt3a, Tet3 and Park2. Exact chromosomal positions of DMCs are given on the x-axis and the orientation of the genes is indicated by an arrow. Hypermethylated are depicted as red squares while hypomethylated DMCs are depicted in blue squares. DMCs were defined as having a methylation difference > or equal to 25 (dashed line) and a q-value <0.01.

## Discussion

Epigenetic alterations and chromatin relaxation are hallmarks of aging ^2, 3^, but evidence of age- and disease-related global changes in the epigenetic landscape of different neuronal populations is still scarce. This information, however, holds important promises for understanding the implication of aging in the neuronal degeneration characterizing age-related NDs and might thus foster the understanding of the pathogenesis of NDs.

Here, we show that *Gadd45b* overexpression induces early changes in DNA methylation, particularly in introns of genes related to neuronal functions and on young and potentially active LINE-1 elements. This, accompanied by perturbations of heterochromatin organization and increased DNA damage, culminate in neuronal cell death after several weeks. At an early time point, before the onset of neurodegeneration, *Gadd45b* overexpression induces a vulnerable state in mDA neurons, increasing their sensitivity to oxidative stress induced by the striatal injection of 6-OHDA. This vulnerability is characterized by an amplified striatal dopaminergic axon terminal loss and an accentuation of perturbations in the organization of heterochromatin and DNA methylation in mDA cell bodies of the SNpc. We also demonstrate that LINE-1 transcripts are increased early on after *Gadd45b* overexpression. This increase in LINE-1, which are potent inducers of DNA damage in mDA neurons ^18^, could explain the cell death of mDA neurons we observe when overexpressing *Gadd45b* long-term.

A number of recent studies indicate that GADD45 proteins play a key role in active DNA demethylation in post-mitotic neurons in the brain ^27, 28^ by serving as scaffolding proteins to recruit DNA repair enzymes such as the thymidine DNA glycosylase (TDG) to the site of DNA demethylation ^44^. So far, GADD45B-regulated DNA demethylation has been described in the context of adult neurogenesis ^29^, depressive-like behavior in mice ^45^, major psychosis in humans ^46^ and cerebral cortex plasticity ^47^ on specific promoters, mostly on promoters of the *Bdnf* gene. Unexpectedly, when overexpressing *Gadd45b* in the SNpc of wild-type mice, we observed very limited changes in methylation on gene promoters, but rather widespread methylation changes on gene bodies with both, hypermethylated and hypomethylated CpGs. This apparent discrepancy with previous studies in terms of extent, localization and pattern of the DNA methylation changes induced by GADD45B might be due to methodology. Indeed, most studies in the brain did not use techniques allowing for unbiased global DNA methylation surveillance.

GADD45B has been shown to associate with TET (Ten-eleven translocation) proteins ^48, 49^ which transform 5-methylcytosines (5mC) through a series of sequential oxidations. Those modified cytosines are excised by TDG and then replaced by non-methylated cytosines through BER-dependent mechanisms ^50^. A recent study has shown that *Tet2* expression is increased in PD patients leading to altered 5mC patterns in enhancers of neuronal genes. Conversely, TET2 loss in mDA neurons was neuroprotective ^51^. *Tet2* did not show any changes in expression at either time point after AAV8-*mGadd45b* injection. This might be explained by the fact that *Tet2* has many splice variants, which were not all covered in our RT-qPCR assay. However, *Tet3* is down-regulated 90d after AAV8-*mGadd45b* injection. TET3 has been shown to interact with transcriptional regulators and histone writers such as H3K36 methyltransferases and to allow the active transcription of certain neuronal genes ^52^. TET3 has also been reported to bind DNA and prevent aberrant methylation at the transcription start site of genes involved in lysosomal functions, mRNA processing and the BER pathway, pointing to a possible relevance of TET3 in the pathogenesis of NDs ^53^. While our knowledge of how genome-wide DNA methylation patterns or epigenetic changes are correlated with PD pathogenesis is still scarce, there is evidence supporting that methylation changes in the SNpc correlate with aging ^23^ and with cognitive impairment in PD ^54^. Most of the existing data concerns methylation patterns of specific genes associated with NDs. Expression of the *SNCA* gene, encoding α-synuclein and mutated in familial PD, for example, is under the control of DNA methylation ^55,56^. Chromatin modifications on specific genes have also been reported, and the expression of several genes mutated in PD are modulated by histone modifications, including *SNCA* and *MAPT* encoding Tau ^25^. Differentially methylated enhancers have been reported in AD patients, and a dysregulation of histone acetylation in both, AD and PD patients ^57–60^.

By serving as an adaptor between repair factors and chromatin, GADD45B can be seen as a communicating platform between DNA repair and epigenetics ^28, 61^. This could provide an explanation to the fact that we observe global changes in heterochromatin organization in addition to changes in DNA methylation. In line with this, GADD45 proteins have been reported to induce heterochromatin relaxation during cellular reprogramming ^31^. GADD45B might thus be part of protein coordinators which link DNA methylation and histone modifications ^60^. Perturbations of DNA methylation and global changes in heterochromatin organization, as we show here, induce neurodegeneration of mDA neurons in the SNpc with time and might thus be primary drivers of neurodegeneration.

Another primary hallmark of aging is genomic instability and some data suggest that this is also true in neuronal aging ^22^. In this context it is important to note, that DNA strand breaks are physiologically occurring in post-mitotic neurons. This physiological process, however, needs to be tightly regulated, since the induction of DNA strand breaks is pathologically exacerbated in AD ^62^. We have shown earlier, that DNA damage in mDA neurons can be induced by acute and chronic oxidative stress ^18, 30^ and that LINE-1 activity participates in DNA damage ^18^. In the latter study, DNA damage was prevented either by siRNA against LINE-1 ORF2, the LINE-1 repressive protein Piwil1 or a nucleoside analogue reverse transcriptase inhibitor *in vitro* ^18^. This LINE-1 induced DNA damage is dependent on young and active LINE-1 copies. These young LINE-1 elements belong mainly to the L1Md-Tf/Gf, A and F families. In this context it is interesting to note that we observe the most pronounced changes in DNA methylation upon *Gadd45b* overexpression on these young LINE-1 families (***Fig. 5***). Intronic DMCs and DMRs, intronic L1-associated DMCs (L1-iDMCs), and DMCs or DMRs overlapping with full-length LINE-1 (flL1-iDMCs/flL1-iDMRs), are majorly hypomethylated and iDMCs are preferentially located in genes related to neuronal functions. It is therefore conceivable that the changes in methylation patterns we observe upon *Gadd45b* overexpression, particularly on young LINE-1 elements, are functionally linked to the increase in LINE-1 transcripts we observe 14 days after injection of AAV8-*mGadd45b*. Furthermore, based on previous evidence summarized above, this increase in expression of young L1 elements of the L1Md-Tf/Gf, A and F families might be at the origin of the DNA damage in mDA neurons upon *Gadd45b* overexpression.

Several lines of evidence suggest that the activation of TEs might be associated with aging and NDs. The expression of LINE-1 in wild-type mice increases in neurons ^63^, liver and muscle during aging ^64^. The activation of TEs with aging leads to neuronal decline and shorter lifespan in drosophila ^65, 66^ and increased TE expression has been reported in brain tissue from PD, AD and ALS patients ^37, 67^. Recently, elevated transcripts of repetitive sequences have been found in the blood of PD patients ^68^. Heterochromatin de-structuration and increased TE activity lead to an AD-like phenotype in a mouse model with targeted disruption of *Bmi1,* a gene involved in heterochromatin maintenance and altered expression in AD patients ^39^. In AD and in ALS, the Tau protein as well as TDP-43 can induce heterochromatin relaxation, especially at the level of LINE-1 elements in the case of TDP-43, leading to increased TE activity and neurotoxicity ^37,67,69^. An aging-induced activation of TEs, which are intrinsic components of the genomes of virtually all eukaryotes, might link genomic instability and epigenetic changes to the aging process.

Changes in chromatin states lead to changes in gene expression as exemplified during the transition from neuronal progenitors to adult neurons ^70^ and during aging ^71^, and recent data suggests that this could also be the case in NDs ^59, 72^. In several NDs (AD and HD ^73, 74^; ALS ^75^), disease-specific gene expression changes are increasingly recognized, but whether they overlap with expression profiles characteristic to aging is not known yet. In AD patients, widespread loss of heterochromatin was accompanied by a transcriptomic profile resembling the one of a fetal brain ^38^, suggesting that the relaxation of chromatin allows the expression of normally repressed genes which alters several biological processes and leads to neurodegeneration ^76^. This is also in line with our data showing the preferential location of iDMCs and iDMRs in genes involved in neurogenesis but it remains to be seen whether their expression is altered. Interestingly, it has been shown that *Gadd45b* activity promotes adult neurogenesis ^29^. The majority of these genes with iDMCs and iDMRs upon overexpression of *Gadd45b* belonged to neuronal categories, particularly synapse-related and neurodevelopmental categories. In this context it is interesting to note that synaptic homeostasis is an emerging key player in the pathogenesis of PD ^77^ and synaptic dysfunction is an early event in neurodegeneration ^78^. *Gadd45b* overexpression induced DMCs and DMRs preferentially in introns of genes. The role of gene body or intronic methylation on gene expression is not completely understood ^79^. In recent years, several studies have reported that gene body methylation influences gene expression levels and/or alternative splicing ^80 81 82^. Among the list of genes underlying a direct or indirect regulation through methylation changes triggered by *Gadd45b* overexpression, the decrease in expression did not correlate with a specific direction of methylation changes towards hypo- or hypermethylated CpGs but rather with a change in the methylation state throughout the gene body. *Satb1* has been described as a dopaminergic-specific regulator of senescence ^83^, a dopaminergic neuron cell survival factor ^84^ and a regulator of global chromatin structuration ^85, 86^. The decline in *Satb1* expression at 90d p.i. might help to explain, at least partly, the neurodegeneration of mDA neurons upon *Gadd45b* overexpression. *Setdb1* is a histone-lysine-methyltransferase that specifically trimethylates lysine-9 of histone H3. It is tempting to speculate that the decrease in expression might relate to the change in the organization of H3K9me3 we observe upon *Gadd45b* overexpression. *Dnmt3a,* a genome-wide *de novo* DNA methyltrasferase, and *Tet3,* are involved in DNA methylation or demethylation, respectively. Interestingly, inactivation of TET and/or DNMT proteins causes gains and losses of DNA methylation, suggesting that the loss of one regulator can lead to the redistribution of other regulators and of DNA modifications ^87^. *Park2,* encoding the ubiquitin protein ligase Parkin, is a PD-related gene and loss-of-function mutations in this gene are responsible for familial forms of PD. It is thus possible, that a decrease in the expression of *Park2* in the context of *Gadd45b* overexpression might contribute to the degeneration of mDA neurons. Overall, the expression changes might be due to the *Gadd45b*-induced epigenetic dysregulation of the neuronal genome and participate in the *Gadd45b* overexpression phenotype and the challenge will be to correlate one with the other.

Altogether, our data are in line with an emerging concept on a new pathogenic pathway initiating age-related neurodegeneration. Recent evidence, including from our group, suggests that aging-induced chromatin reorganization triggers the activation of LINE-1 retrotransposons and subsequent LINE-1 induced DNA damage cumulating in neuronal cell death. Our group has shown that acute and chronic oxidative stress leads to heterochromatin relaxation and LINE-1 activation in mDA neurons *in vivo* ^18, 30^. Aging-induced epigenetic alterations might produce a vulnerable pre-ND state. Combining this pre-ND state with a particular genetic susceptibility, a familial gene mutation or an accelerating environmental trigger, could initiate a cascade of secondary events including protein aggregation, metabolic dysregulation and mitochondrial dysfunctions. NDs share several common pathological features and despite extensive investigation, no disease-modifying treatment is available. Acknowledging aging as a vulnerability factor for neurodegeneration is important, not only for understanding the pathogenesis of NDs, but also for modeling, testing and developing therapeutics for crucially lacking disease-modifying treatments. Our study suggests two novel therapeutic targets for neuroprotection. Drugs restoring chromatin structure and/or repressing LINE-1 transcription or activity might hold promise for the prevention of age-related neurodegeneration.

## Materials and Methods

### Animals

All animal treatments followed the guidelines for the care and use of laboratory animals (US National Institutes of Health), the European Directive number 2010/68/UE (EEC Council for Animal Protection in Experimental Research and Other Scientific Utilization). This project was validated by the competent ethical committee (CEA 59) and authorized by the Minister of Higher Education, Research and Innovation (n° 00703.01 and n° APAFIS#6605-2016090209434306 v3). For surgical procedures, animals were anesthetized with Xylazine (Rompun 2%, 5 mg/kg) and Ketamine (Imalgene 1000, 80 mg/kg) by intraperitoneal injection and a local subcutaneous injection of lidocaine (0.5%, 3mg/kg) on the incision site. Post-chirurgical analgesia was insured by an injection of the analgesic Meloxicam (Metacam, 0,5mg/kg) s.c.. Swiss OF1 wild-type mice (Janvier) were maintained under a 12 h day/night cycle with ad libitum access to food and water. A maximum of 5 mice were housed in one cage, and cotton material was provided for mice to build a nest. Experimental groups consisted of five to seven male mice of 6 weeks of age. Sample size calculations were based on previous experiments.

### AAV8 vectors to overexpress Gadd45b

Forced expression of *Gadd45b* in mDA neurons was achieved using an AAV8 viral vector. The constructs contained cDNAs for either mouse *Gadd45b* (AAV8-*mGadd45b*) or *mCherry* (AAV8-*mCherry*) under the control of the ubiquitous EF1a promotor. *Gadd45b* cDNA was flanked by the cognate 5’ and 3’ UTRs (***Fig. 1A***). AAV8 was chosen because it has previously been shown to efficiently infect mDA neurons in the midbrain.

### Brain injections

For injections, mice were placed in a stereotaxic instrument, and a burr hole was drilled into the skull 3.3 mm caudal and 1.3 mm lateral to the Bregma. The needle was lowered 3.8 mm from the surface of the skull, and AAV8-Ef1a-*mCherry* or AAV8-Ef1a-*mGadd45b* (Vector Biolabs; 2 μl; 4.8×10^13^ GC/ml suspended in NaCl 0,9% with 5% glycerol) injections were performed over 4 min. Where indicated 6-OHDA (2 μl; 0.5 gg/gl; Sigma) injections were performed in the same manner 0.4 mm rostral, 1.8 mm lateral and 3.8 mm ventral to the bregma, over 4 min.

### Tissue dissection

For RNA and DNA analyses, biopsies of the SNpc were performed. Brains were put into a custom-made brain slicer for adult mice brain. A coronal slice of ≈2mm encompassing the SNpc was excised (Bregma −3.26 mm to −5.2mm) and placed on a cold cover slide with the rostral side facing the experimenter. Dissection of the SNpc was then done following anatomical landmarks: a sagittal cut to separate the two hemisphere, a second parasagittal cut through the fasciculus retroflexus and the mammillothalamic tract (about 2/3 starting from the midline of the distance between the midline and the rostral end of cerebral peduncle) to remove the VTA, a transversal section from the ventral part of the lateral geniculate complex to the midline, a second transversal cut from the ventral end of the cerebral peduncle to the midline. The cerebral cortex was then removed to only keep the midbrain part containing the SNpc and immediately frozen on dry ice and kept at −80°C until extraction.

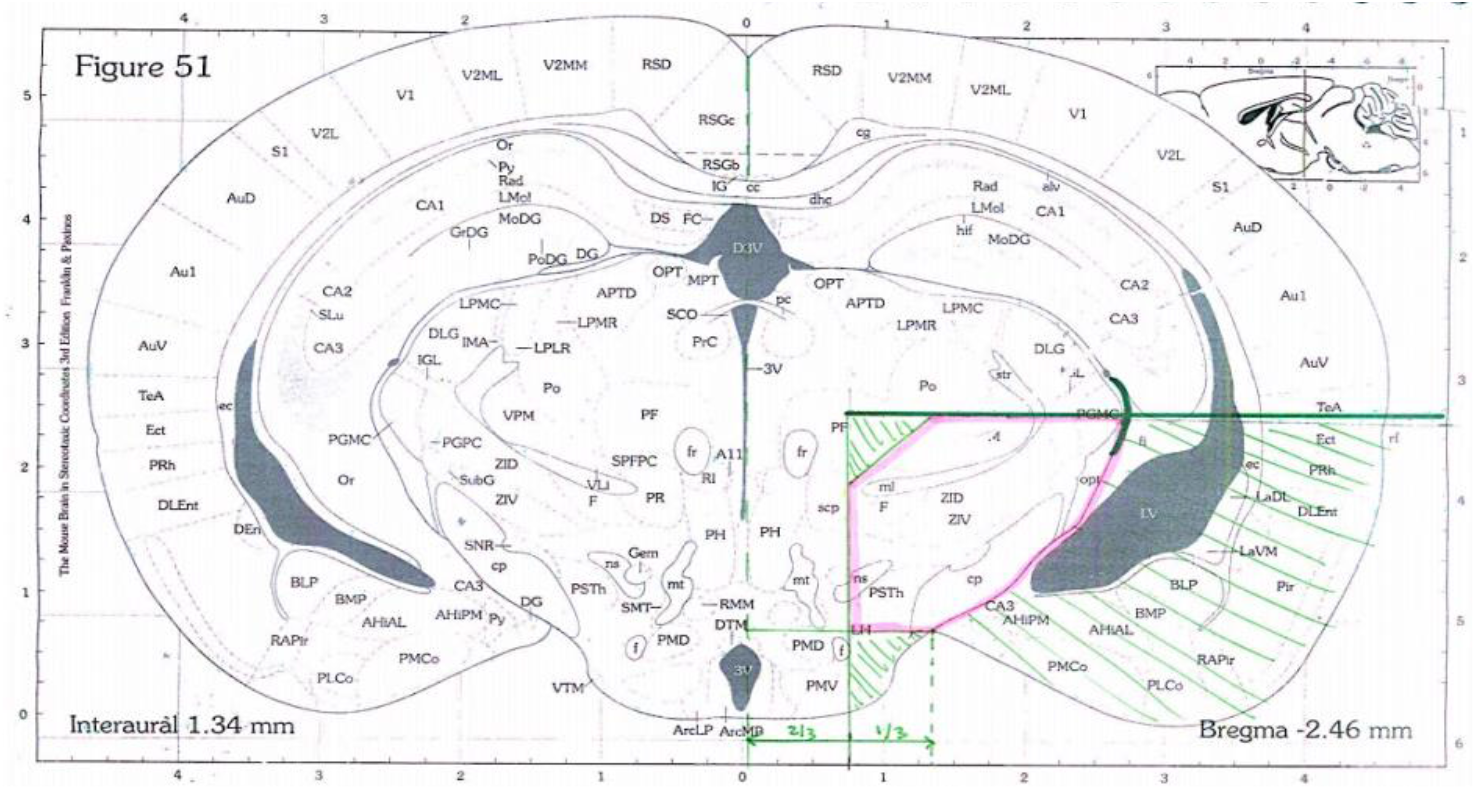

### RT-qPCR

Total RNA from dissected SNpc was extracted using the AllPrep DNA/RNA Micro Kit (Qiagen 80284) adding an on-column DNase I treatment (Qiagen 79256), followed by RT-qPCR. RNA (200 ng) was reverse-transcribed using the QuantiTect Reverse Transcription kit (Qiagen 205313). Quantitative PCR reactions were carried out in duplicates with SYBR Green I Master Mix (Roche S-7563) on a LightCycler 480 system (Roche Applied Science). The following primers were used: Hprt (sense: AGCAGGTGTTCTAGTCCTGTGG, antisense: ACGCAGCAACTGACATTTCTAA); LINE-1 A (sense: TTCTGCCAGGAGTCTGGTTC, antisense: TGAGCAGACCTGGAGGGTAG); LINE-1 Tf/Gf (sense: CTGGGAACTGCCAAAGCAAC, antisense: CCTCCGTTTACCTTTCGCCA); Gadd45b (sense: ACTGATGAATGTGGACCCCG, antisense: CCTCTGCATGCCTGATACCC); Satb1 (sense: TCTTTTACCCCCTCCTCCCA, antisense: TCACCTGCCAGAACACTTCA); Tet3 (sense: CTCGGCGGGGATAATGGGAG, antisense: AGCCTGTCTTGACAGTCGCC); Dnmt3a (sense: GCCGAATTGTGTCTTGGTGGATGACA, antisense: CCTGGTGGAATGCACTGCAGAAGGA), Setdb1 (sense: GTTTGCCTGGGTTTGGCAAG, antisense: CTTTGGCCCTCAGTCCGTC); Park2 (sense: GCTCAAGGAAGTGGTTGCTAAG, antisense: CAATACTCTGTTGTTCCAGGTCA). Primer efficiencies were tested using 10-fold dilution series of cDNA spanning at least three orders of magnitude. Data were analyzed using the ddCt method and values normalized to hypoxanthine-guanine phosphoribosyl transferase (Hprt).

### DNA extraction and quantification

DNA was also extracted during the same process of RNA extraction using the AllPrep DNA/RNA Micro Kit (Qiagen 80284). The DNA was then purified, treated with RNase H (ThermoFischer, 18021071) and Proteinase K (PCR grade, Roche, 3115836001) and concentrated with the DNA Clean & Concentrator-5 kit (Zymo, D4013). DNA concentration of each sample was measured using the Qubit® fluorometer with dsDNA BR Assay Kit (Thermo Fisher Scientific).

### RRBS

RRBS was performed by Diagenode. DNA quality of the samples was assessed with the Fragment AnalyzerTM and the DNF-488 High Sensitivity genomic DNA Analysis Kit (Agilent). DNA was slightly more fragmented than defined by the quality control standards but this fragmentation was minor. RRBS libraries were prepared using the Premium Reduced Representation Bisulfite Sequencing (RRBS) Kit (Diagenode) which uses the Mspl restriction enzyme and size selection to enrich for CpG-rich regions (coverage of about 4 million CpGs). 100ng of genomic DNA were used to start library preparation for each sample. Following library preparation, samples were pooled together by 8. PCR clean-up after the final library amplification was performed using a 1.45x beads:sample ratio of Agencourt® AMPure® XP (Beckman Coulter). DNA concentration of the pools was measured using the Qubit® dsDNA HS Assay Kit (Thermo Fisher Scientific). The profile of the pools was checked using the High Sensitivity DNA chip for 2100 Bioanalyzer (Agilent). RRBS library pools were sequenced on a HiSeq3000 (Illumina) using 50 bp single-read sequencing (SR50). Bisulfite conversion and amplification were performed using Diagenode’s Premium RRBS Kit. After conversion, the pooled samples were analyzed by qPCR. Sequencing was performed in single-end mode, generating 50 bases reads (SE50) on an Illumina HiSeq 3000/4000. Quality control of sequencing reads was performed using FastQC version 0.11.8 (https://www.bioinformatics.babraham.ac.uk/projects/fastqc/). Adapter removal was performed using Trim Galore (https://www.bioinformatics.babraham.ac.uk/projects/trim_galore/) version 0.4.1. Reads were then aligned to the murine reference genome mm10/GRCm38 using bismark v0.20.0 ^88^. Bismark is a specialized tool for mapping bisulfite-treated reads such as the ones generated in RRBS-seq experiments. Bismark requires that the referenced genome first undergoes an in-silico bisulfite conversion while transforming the genome into forward (C → T) and reverse strand (G → A) versions. The reads producing a unique best hit to one of the bisulfite genomes were then compared to the unconverted genome to identify cytosine contexts (CpG, CHG or CHH - where H is A, C or T). The cytosine2coverage and bismark_methylation_extractor modules of Bismark were used to infer the methylation state of all cytosines (for every single uniquely mappable read) and their context, and to compute the percentage methylation. The reported cytosines were filtered to get only the CpGs covered in each sample. The spike-in control sequences were used at this step to check the bisulfite conversion rates and to validate the efficiency of the bisulfite treatment. Methylkit v1.7.0 ^89^, a R/Bioconductor package, was used to perform the differential methylation analysis between the two groups of samples. The CpG data set was filtered for low coverage (CpGs with coverage less than 10X in all samples per comparative group were discarded) and for extremely high coverage to discard reads with possible PCR bias (CpGs with coverage higher than the 99.9th percentile were discarded). The data was then normalized for read coverage distribution between samples. A pairwise comparison was performed for first versus second group of samples to identify differentially methylated CpGs (DMCs) and differentially methylated regions (DMRs), the latter with a window and step size of 1000bp. Methylkit uses logistic regression to compare the methylation percentages between groups at a given CpG/region. All DMCs and DMRs were annotated with the R/Bioconductor package annotation ^90^, with the refGene and CpG island annotations from UCSC. The annotation comprised two categories: (i) distance to a CpG island and (ii) regional annotation. The distance related annotation classified DMCs and DMRs whether they overlapped a known CpG island, 2000 bp of the flanking regions of the CpG islands (shores), 2000 bp of the flanking regions of the shores (shelves) or outside these regions (open sea). The regional analysis classified DMCs and DMRs in four groups, namely, exons, introns, promoters and intergenic regions.

| Sample Name | Total reads | Uniquely aligned | Mapping efficiency (%) | CpGs detected | CpGs cov>10 | Average Coverage | Conv. Rate Meth. spike-in (%) | Conv. Rate Unmeth. spike-in (%) |
| --- | --- | --- | --- | --- | --- | --- | --- | --- |
| mC1_2w | 32.674.187 | 21.455.193 | 65.66 | 1.552.817 | 1.316.296 | 36.96 | 1.76 | 99.34 |
| mC4_2w | 45.583.765 | 29.506.694 | 64.73 | 1.786.123 | 1.544.593 | 38.73 | 1.56 | 99.53 |
| mG1_2w | 39.606.244 | 26.791.258 | 67.64 | 1.727.650 | 1.495.577 | 40.53 | 1.48 | 99.30 |
| mG6_2w | 43.911.896 | 29.422.323 | 67.00 | 1.742.579 | 1.527.329 | 43.18 | 1.56 | 99.47 |
| mC1_3m | 43.426.559 | 28.584.928 | 65.82 | 1.719.287 | 1.499.415 | 40.19 | 1.31 | 99.51 |
| mC3_3m | 37.983.254 | 24.904.215 | 65.57 | 1.644.193 | 1.412.632 | 37.57 | 1.64 | 99.42 |
| mG1_3m | 41.810.446 | 27.906.101 | 66.74 | 1.665.705 | 1.481.471 | 44.30 | 1.73 | 99.52 |
| mG4_3m | 37.699.685 | 24.957.045 | 66.20 | 1.652.248 | 1.422.694 | 36.72 | 1.72 | 99.47 |

### Immunostaining

For immunostaining, animals received a lethal intraperitoneal injection of 1 μl/g body weight dose of Euthasol (150mg/kg) and were then perfused with 8 mL of Phosphate Buffer Saline (PBS) then 8 mL of 4% Paraformaldehyde (PFA) at a rate of 300 ml/h using a syringe pump. Brains were then post-fixated 1 hour at room temperature (RT) in 4% PFA, washed in PBS three times for 30 minutes at RT and placed in PBS with 20% sucrose overnight at 4°C. After cryoprotection, the brains were embedded in Tissue Freezing Medium (TFM, Microm Microtech), frozen on dry ice and 30 μm sections of mouse striatum and ventral midbrains encompassing the SNpc were prepared using an HM 560 Microm cryostat (Thermo Scientific).

Slides with 30 μm striatum or midbrain sections were washed in PBS and permeabilized with 1% Triton X-100. After 30 minutes at RT in 100 μM glycine buffer (for TH/mCherry and TH/ORF1p) or 30 minutes at 100°C in demasking citrate buffer (10 mM, pH 6, 0.05% Tween) (for TH/MeCP2, H3K9 or γ-H2AX), sections were first blocked in 10% Fetal Bovine Serum (FBS, Gibco) in the presence of 0.5% Triton X-100 for 1 hour at RT and incubated with primary antibodies overnight at 4°C, washed and further incubated with secondary antibodies for 1 hour at RT. The following primary antibodies used: anti-γ-H2AX (mouse, 1/200, Millipore, clone JBW301), anti-TH (chicken, 1/500, Abcam, ab76442), anti-ORF1p (guinea pig, 1/200, in-house, clone 09 as in ^83^, anti-mCherry (mouse, 1/200, Clontech 632543), rabbit anti-H3K9me3 (rabbit, 1/200, Abcam, ab8898) and anti-MeCP2 (rabbit, 1/200, Millipore MABE328). Sections were incubated with appropriate secondary antibodies (488 anti-chicken, 546 anti-mouse, 647 anti-guinea pig, 647 anti-rabbit, 647 anti-mouse, Alexa Fluor, Life Technologies)

### In situ hybridization

Mice were perfused with PBS in RNase-free conditions, and frozen in isopentane (embedded in TissueTek O.C.T). Brain slices (20 μm) were fixed in 4% PFA in PBS for 10 min at RT and then permeabilized twice for 10 min in RIPA buffer (150 mM NaCl, 1% NP-40, 0.5% Na deoxycholate, 0.1% SDS, 1 mM EDTA, 50 mM Tris–HCl pH 8). Brain sections were fixed again for 5 min in 4% PFA, acetylated for 10 min in 0.25% acetic anhydride (in 0. 1 M triethanolamine, pH 8). Sections were then permeabilized for 30 min in PBS with 1% Triton X-100, 10 min in 10mM citrate buffer pH6 at a 100°C and then pre-incubated for 1 h at 70°C in hybridization buffer (50% formamide, 5× SSC, 5× Denhardt (1% Ficoll, 1% SSC, 1% Tween-20), 500 μg/ml Salmon sperm DNA, 250 μg/ml yeast tRNA). Sections were then hybridized overnight at 70 °C with a digoxigenin (DIG)-labeled RNA probes (DIG RNA labeling kit, Roche 11277073910) for *Gadd45b* mRNA. Sections were washed with FAM/SSC (50% formamide, 2× SSC, 0.1% Tween-20) twice 30 min at 37°C, then twice in 0.2× SCC at 42°C, blocked in B1 buffer (100 mM maleic acid pH 7.5, 150 mM NaCl) with 10% fetal bovine serum (FBS) for 1 h, and incubated overnight at 4°C in B1 buffer with alkaline phosphatase-conjugated anti-DIG (1/2000; Roche 11633716001). After three washes in B1 buffer and one wash in B3 buffer (100 mM Tris–HCl pH 9, 50 mM MgCl2, 100 mM NaCl, 0.1% Tween-20) for 15 min, slides were stained using the NBT/ BCIP kit (Vector labs, SK5400), stopped with ddH20 and slides were mounted with DAKO-mounting medium.

### Imaging/ Microscopy

All large field images used for TH+ neuron quantification, level of viral infection quantification and striatal intensity quantification were made on widefield microscope (Axio zoom V16 – Zeiss – Apotome.2) at 2,3 magnification with a zoom factor of 100. ISH image was taken by upright widefield microscope equipped with a color CCD camera (Nikon 90i microscope) at 20x magnification in brightfield. H3K9me3, MeCP2 and γ-H2AX foci quantification as well as ORF1p intensity quantification were made on images taken by spinning disk microscopy (Yokogawa W1 Spinnning-disk head mounted on an inverted Zeiss AxioObserver Z1) at 63x magnification.

The images in the figures are for illustration purposes and were taken by confocal microscopy (LSM 980 with Airyscan 2, Zeiss) at 63x magnification with a zoom factor of 1.4, except for the MeCP2 and ORF1p images (performed on the spinning-disk microscope).

### Cell counting and image quantification

TH cell counting in conditions comparing ipsi- (treated) and contralateral (non-treated) sides were done as follows: For every brain, a minimum of four serial sections were stained, and the number of TH cells was counted in the SNpc of both ipsi- and contralateral sides. An ipsi/contra ratio was calculated for each section, and the resulting mean of four sections was used to quantify the difference between the TH cell number of the ipsi- and contralateral side of the same animal. The counting was done blindly.

The quantification of axonal degeneration in the striatum comparing ipsi- and contralateral sides was done as follows: For every brain, a minimum of seven serial sections were stained, and the integrated density of TH staining intensity was measured in ImageJ by determining the entire contralateral striatum as region of interest (ROI) and conserving area for the measurement of ipsilateral sides. An ipsi/contra ratio was calculated for each section, and the mean ratio of sections containing the striatum was used to quantify the difference between TH striatal intensity of the ipsi- and contralateral side of the same animal. The quantification was done blindly.

Quantifications of foci were performed using a 63× magnification and 0.3 μm - thick successive focal planes except for γ-H2AX foci quantification, which was made using 0.2 μm-thick successive focal planes. Immunostainings of one parameter (H3K9, MeCP2, Etc.) were all done in one experiment for all the conditions. For each immunostaining, 3 images of the SNpc per side were taken on 3 sections per mouse, thus 18 images per mouse (n=3 per condition), or 54 images per condition. The same parameters were set-up on the spinning disk microscope to allow for comparison between experimental conditions for the same staining. Images were analyzed by the same experimenter using ImageJ software ^84^. For foci quantifications the foci counting Fiji Plug-in was used ^85^. In addition, an image analysis plug-in was developed for the ImageJ/Fiji software, using Bio-Format (openmicroscopy.org), mcib3D ^86^ and GDSC (Alex Herbert from Sussex University) libraries. First, nuclei that belonged to TH+ neurons were manually marked with the plug-in Cell Counter, an xml file for each image containing 3D nuclei coordinates was saved. Then, nucleus channel was filtered with a median filter (radius = 4) and a Difference of Gaussian (DOG) (sigma1 = 30, sigma2 = 15), a binary mask was done with an Otsu threshold. Only nuclei that are associated to the nuclei defined in the xml file were kept. MeCP2 and H3K9me3 foci detections were performed using a median filter (radius=2), DOG (sigma1 =10, sigma2=2), binary mask was done with a MaxEntropy threshold, then 3D objects (foci) that had a volume comprised between 1.5 and 40 μm^3^ and were inside TH+ nuclei or at a 2 μm distance to nucleus was associated to nuclei. γ-H2AX foci detection was performed using a median filter (radius=2), DOG (sigma1=7, sigma2=3), binary mask was done with a Moments threshold, then 3D objects (foci) that had a volume comprised between 0.5 and 100 μm^3^ were inside TH+ nuclei or at a 2 μm distance to nucleus was associated to nuclei. For each nucleus, foci number, average foci volume, average foci integrated intensity and average nucleus integrated intensity was computed.

### Gene ontology analysis

Gene ontology analysis (PANTHER version 15.0; ^87^) was done using the PANTHER overrepresentation test with the GO-Slim annotation data sets ‘biological process’, ‘molecular function’ and cellular component’ and the *‘mus musculus’* gene set as the reference list. Fisher’s exact test was used to compute statistical significance of overrepresentation with the false discovery rate (FDR) set at p < 0.05. The first 15 categories, ordered by fold enrichment, are displayed along with the corresponding FDR value.

### Statistics

Unless otherwise stated, the graphs represent each replicate and the error bar the SEM of the mean of replicates. Error bars, values of n and mean ± SEM are as stated in the figure legends. Results were considered as statistically significant for P-value <0.05; in some cases, the exact P-value is given. Normality test were performed prior to the statistical test and unless stated otherwise, the nonparametric Wilcoxon-Mann-Whitney test was applied. All statistical analysis was done with the software Prism.

For the bioinformatic analysis of the RRBS, we formulated the null hypothesis that there are no differences in methylation between the two groups. After the p-values have been computed, Methylkit, an R package for DNA methylation analysis and annotation, uses the sliding window model (SLIM) to correct P-values to q-values for multiple comparison tests. Statistically significant DMCs and DMRs were identified with a q-value cutoff <0.01 and a methylation difference higher than 25%.

## Acknowledgements

This work was supported by the Fondation de France (00086320 to J.F.) and the Fondation du Collège de France (to J.F.). We thank all primary donors for their financial contributions to this work. We thank the animal facility members for their essential contributions. We gratefully acknowledge Julien Dumont and the Collège de France Orion imaging facility (IMACHEM-IBiSA), member of the French National Research Infrastructure France-BioImaging (ANR-10-INBS-04), which received support from the program «Investissements d’Avenir» ANR-10-LABX-54 MEMOLIFE. We thank Yves Dupraz for the manufacturing of a customized mouse brain slicer.

## Author contributions

*CRG performed most of the experiments and participated in the writing of the manuscript, OMB contributed experimentally, PM designed the semi-automated image analysis workflow, AP co-supervised the beginning of the study, RLJ co-supervised the study and contributed to the writing of the manuscript, JF co-supervised the study, analyzed the RRBS data, wrote the manuscript and received the funding.*

