## Supplementary data for "Perturbed DNA methylation by sustained overexpression of Gadd45b induces chromatin disorganization, DNA strand breaks and dopaminergic neuron death in mice"

***Supplementary Figures***

***Suppl. Figure 1. Gadd45b overexpression in the SNpc of wild-type mice leading to DNA methylation changes.***

***A, C:*** *Volcano plots of differentially methylated regions.*

*Volcano plots compare Gadd45b versus mCherry 14d p.i. (**A**) and 90d (**C**). 16 702 (14d p.i.) and 15 685 (90d p.i.) significantly differentially methylated regions were detected with a q-value smaller than, or equal to, 0.01 and at least 25% difference. The Volcano plot shows the number of regions with changed patterns of methylation, between Gadd45b and control samples, which are significantly higher or lower than the 25% difference cut-off and considering a q- value threshold of 0.01. The difference in methylation is reflected in the x-axis while the y-axis represents the significance of the difference. Regions that are highly differentially methylated are further to the left and right sides of the plot, while highly significant changes appear higher on the plot. Values on the x- and y-axes are percent methylation differences and negative log10 of the corrected p-values, respectively. The pie chart shows the percentage of hyper and hypo methylated regions.*

***B, D****: Annotation of DMRs.*

*DMRs were annotated in relation to the distance to a CpG island 14d p.i. (**B, left panel**) and 90d p.i. (**D, left panel**), as well as based on the genomic regions they are associated with 14d p.i. (**B, right panel**) and 90d p.i. (**D, right panel**). Their distributions are plotted in a bar chart.*

***E:*** *Venn diagrams of overlapping genes with intronic DMRs. Venn diagrams displaying the overlap of genes containing at least one hypomethylated (left) or hypermethylated intronic DMR (right) at 14d and 90d p.i.. The overlap of genes with hyper- or hypomethylated intronic DMRs that are conserved between 14d and 90d revealed 447 genes containing at least one intronic hypermethylated as well as at least one hypomethylated DMR at both 14d and 90d.*

***F:*** *Gene ontology analysis of the "Gadd45b-DMR-regulon". The PANTHER overrepresentation test with the GO-Slim annotation data set 'biological process' identified significantly overrepresented GO categories (447 genes). The first fifteen significantly overrepresented GO categories with the highest fold enrichment are displayed with the fold enrichment on the left y-axis (black points) and the FDR value on the right y-axis (grey points).*

**G**: *CpG methylation percentage by chromosome for DMCs. The percentage of hypo- and hypermethylated CpGs at 14d p.i. was plotted per chromosome (q-value<0.01; methylation difference > or equal to 25%), hypomethylated CpGs are represented in blue and hypomethylated CpGs in red.*

***Suppl. Figure 2. Analysis of MeCP2 foci in TH+ neurons upon Gadd45b overexpression.***

*There are no significant differences in the number (2.72±0.13; 2.45±0.12), volume (2.98±0.12; 3.27±0.15 $\mu m^3$) or intensity (9.69x$10^5$±36119; 10.16x$10^5$±40031 UA) of MeCP2 foci in TH+ neurons of the SNpc at 14d p.i. although there is a slight 1.07-fold increase in the intensity of the diffuse nucleoplasmic staining (3.47x$10^7$±774853; 3.71x$10^7$±883600 UA) in TH+ neurons of the SNpc of AAV8-mGadd45b mice compared to AAV8-mCherry mice, shown in (**A, B**), quantified in (**E, F, G**). There are also no significant differences in the number (2.08±0.12; 2.13±0.10), volume (4.40±0.23; 4.84±0.23 $\mu m^3$), intensity (1.20x$10^6$±51929; 1.35x$10^6$±53221 UA) of MeCP2 foci or in the diffuse nucleoplasmic intensity of MeCP2 staining (3.59x$10^7$±888270; 3.72x$10^7$±715283 UA) at 90d p.i., in TH+ neurons of the SNpc, shown in (**C**), quantified in (**D, H, I, J**). Scale bar represents 5 $\mu$m; error bars represent SEM; * p<0.05; n=3. Between 485 and 552 neurons at 14d p.i. and between 379 and 512 neurons at 90d p.i. were quantified per condition.*

***Suppl. Figure 3. LINE-1 methylation changes and increased LINE-1 expression following Gadd45b overexpression.***

***A-C**: Analysis of DMRs in LINE-1 sequences (L1-DMRs) at 14d p.i. of AAV8-Gadd45b.*

*Of a total of 1031 DMRs overlapping with an annotated LINE-1 element, 264 (25,6 %) L1-DMRs were located in intronic regions and 766 in intergenic regions (**A**). The majority of LINE-1 associated DMRs located in introns (L1-iDMRs) were hypomethylated (63,3 %, 167 L1-iDMRs, **A**), located in protein-coding genes (83,7 %, 221) and of the L1Md_F family (**B**). Note that hypo- and hypermethylated LINE-1 elements of the active L1Md_T (3 and 2, respectively) and L1Md-A families (3 and 3 elements, respectively) were present.*

***(C):** Overlap of DMRs and DMCs with an annotation of full-length LINE-1 elements (L1Basev2). Ten DMRs and 62 DMCs overlapped with a full-length LINE-1 element, of which in both cases more than half (7 out of 10 DMRs and 38 out of 62 DMCs), located in either introns or intergenic, were hypomethylated. Are depicted here the genes either containing* *an intronic full-length LINE-1 with DMRs or nearest one in the case of intergenic full-length LINE-1 with DMRs (in white boxes).*

***Suppl. Figure 4. Expression of selected genes in the SNpc.***

*RNA-seq data (base mean expression) of RNA extracted from laser capture micro-dissected SNpc of wildtype mice (pooled from n=4 mice each,* GEO *GSE72321;* [30] and *unpublished*) *was interrogated concerning the expression of several dopaminergic markers (TH, Slc6a3 and En1) and compared to the expression levels of Gadd45b and gene candidates.*

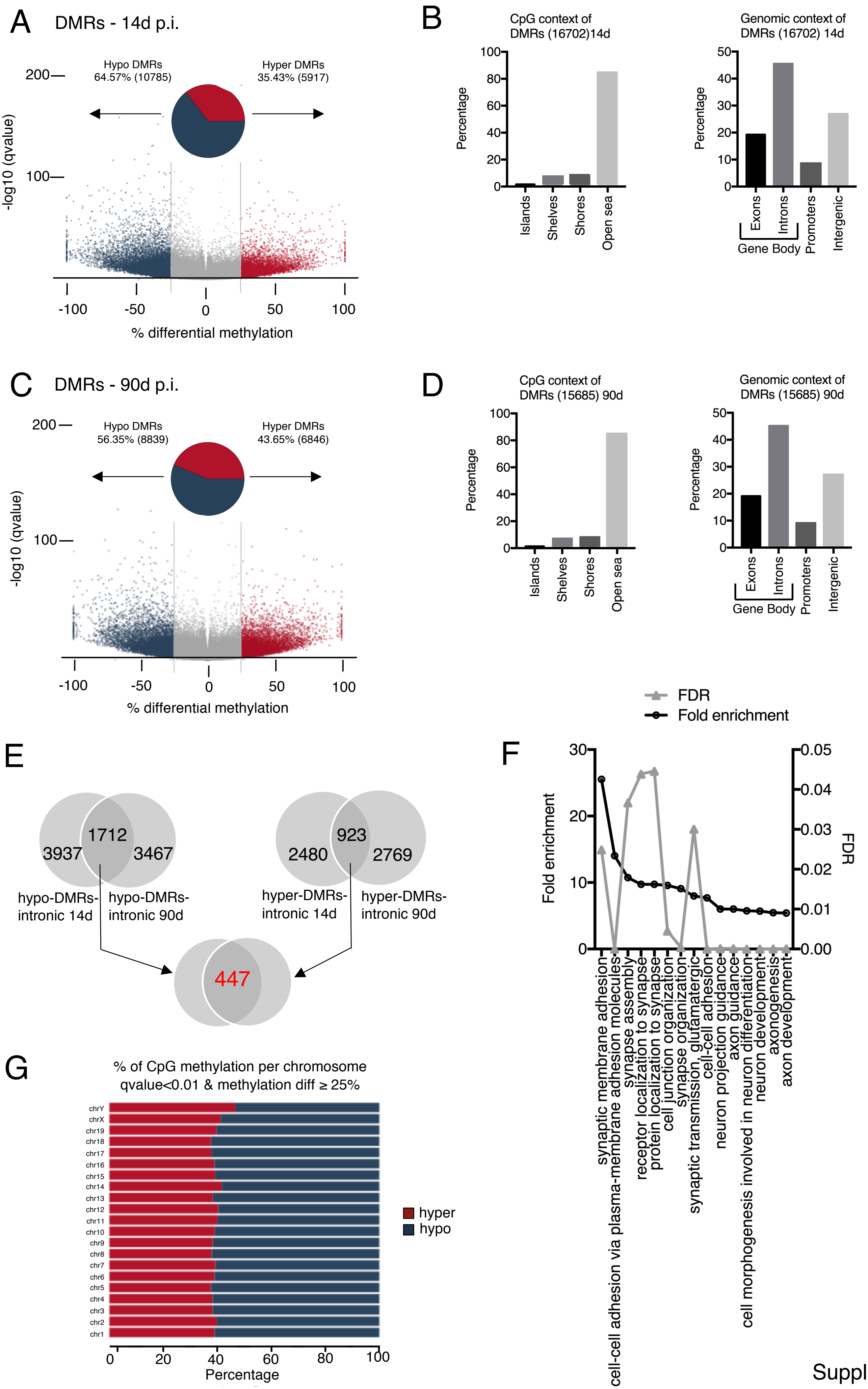

A
DMRs - 14d p.i.
Hypo DMRs 64.57% (10785)
Hyper DMRs 35.43% (5917)
-log10 (qvalue)
% differential methylation
B
CpG context of DMRs (16702)14d
Genomic context of DMRs (16702) 14d
Percentage
Islands
Shelves
Shores
Open sea
Exons
Introns
Gene Body
Promoters
Intergenic
C
DMRs - 90d p.i.
Hypo DMRs 56.35% (8839)
Hyper DMRs 43.65% (6846)
-log10 (qvalue)
% differential methylation
D
CpG context of DMRs (15685) 90d
Genomic context of DMRs (15685) 90d
E
1712
3937
3467
hypo-DMRs-intronic 14d
hypo-DMRs-intronic 90d
923
2480
2769
hyper-DMRs-intronic 14d
hyper-DMRs-intronic 90d
447
F
FDR
Fold enrichment
synaptic membrane adhesion
cell-cell adhesion via plasma-membrane adhesion molecules
synapse assembly
receptor localization to synapse
protein localization to synapse
cell junction organization
synapse organization
synaptic transmission, glutamatergic
cell-cell adhesion
neuron projection guidance
axon guidance
cell morphogenesis involved in neuron differentiation
neuron development
axonogenesis
axon development
G
% of CpG methylation per chromosome
qvalue<0.01 & methylation diff ≥ 25%
hyper
hypo
Percentage


Suppl. Fig. 1

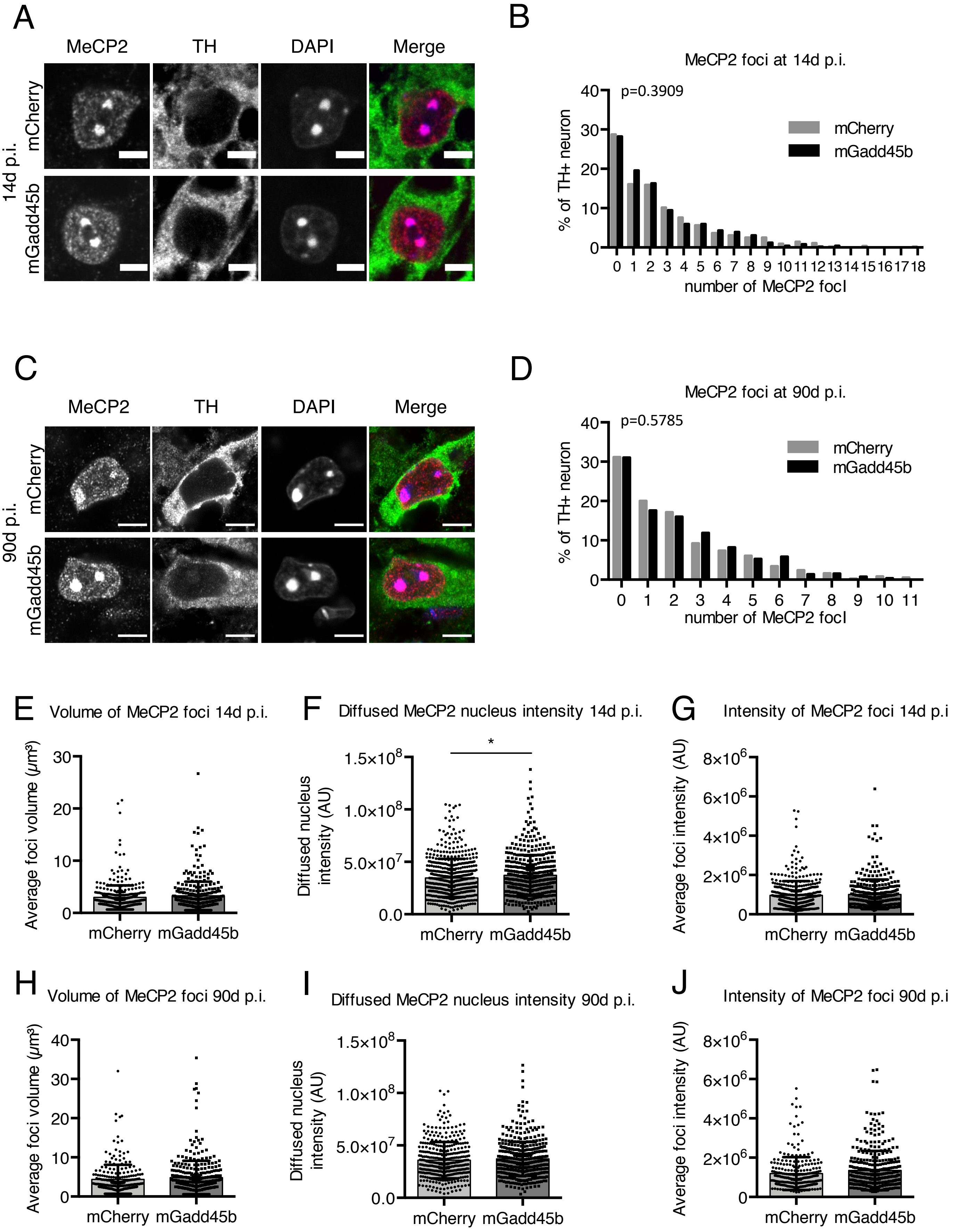
A
MeCP2
TH
DAPI
Merge
14d p.i.
mCherry
mGadd45b
B
MeCP2 foci at 14d p.i.
p=0.3909
mCherry
mGadd45b
% of TH+ neuron
number of MeCP2 focI
C
MeCP2
TH
DAPI
Merge
90d p.i.
mCherry
mGadd45b
D
MeCP2 foci at 90d p.i.
p=0.5785
mCherry
mGadd45b
% of TH+ neuron
number of MeCP2 focI
E
Volume of MeCP2 foci 14d p.i.
Average foci volume (μm³)
mCherry mGadd45b
F
Diffused MeCP2 nucleus intensity 14d p.i.
Diffused nucleus intensity (AU)
*
mCherry mGadd45b
G
Intensity of MeCP2 foci 14d p.i
Average foci intensity (AU)
mCherry mGadd45b
H
Volume of MeCP2 foci 90d p.i.
Average foci volume (μm³)
mCherry mGadd45b
I
Diffused MeCP2 nucleus intensity 90d p.i.
Diffused nucleus intensity (AU)
mCherry mGadd45b
J
Intensity of MeCP2 foci 90d p.i
Average foci intensity (AU)
mCherry mGadd45b

Suppl. Fig. 2

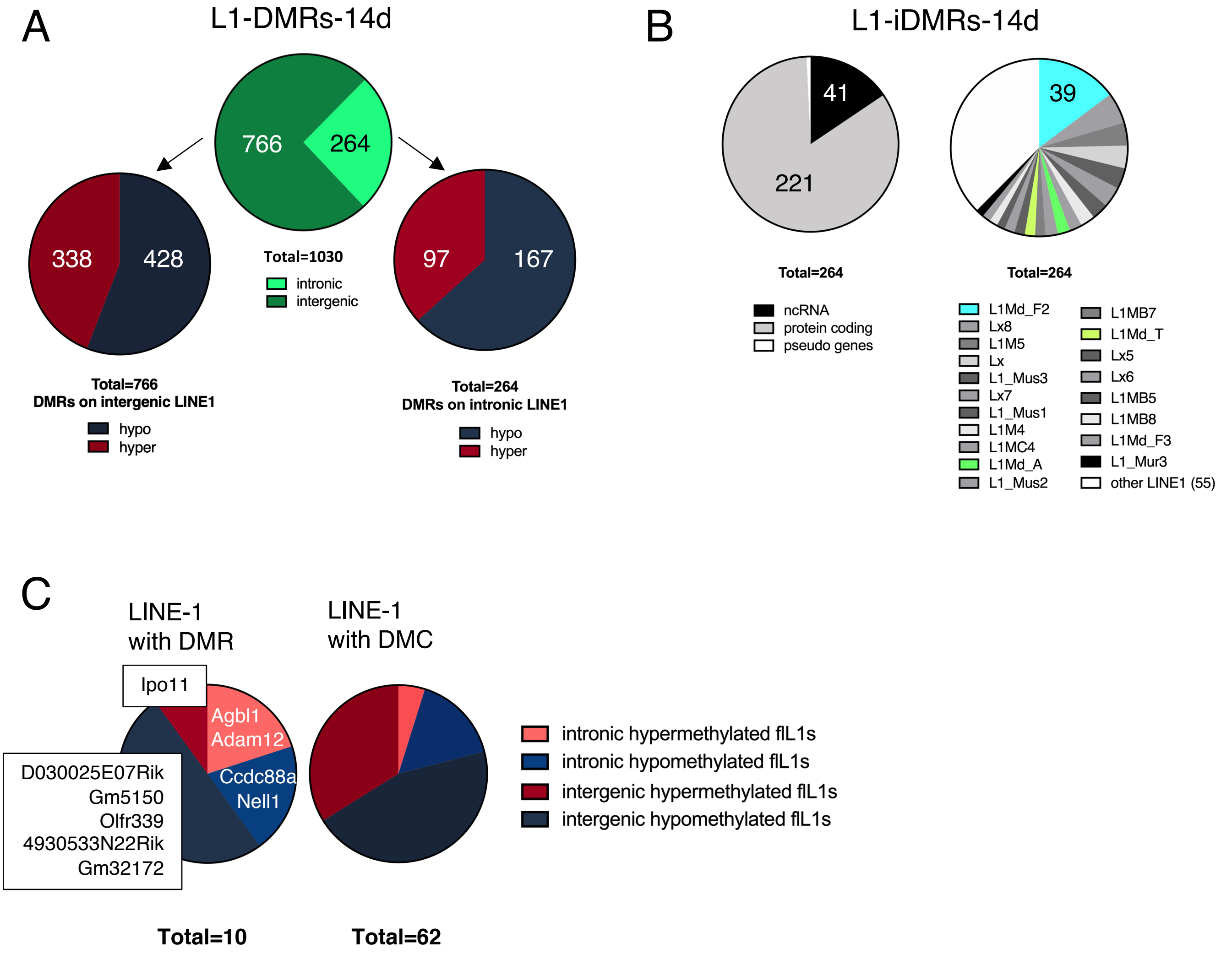
A
L1-DMRs-14d
766
264
Total=1030
intronic
intergenic
338
428
Total=766
DMRs on intergenic LINE1
hypo
hyper
97
167
Total=264
DMRs on intronic LINE1
hypo
hyper
B
L1-iDMRs-14d
41
221
Total=264
ncRNA
protein coding
pseudo genes
39
Total=264
L1Md_F2
Lx8
L1M5
Lx
L1_Mus3
Lx7
L1_Mus1
L1M4
L1MC4
L1Md_A
L1_Mus2
L1MB7
L1Md_T
Lx5
Lx6
L1MB5
L1MB8
L1Md_F3
L1_Mur3
other LINE1 (55)
C
LINE-1
with DMR
Ipo11
Agbl1
Adam12
Ccdc88a
Nell1
D030025E07Rik
Gm5150
Olfr339
4930533N22Rik
Gm32172
LINE-1
with DMC
intronic hypermethylated flL1s
intronic hypomethylated flL1s
intergenic hypermethylated flL1s
intergenic hypomethylated flL1s
Total=10
Total=62

Suppl. Fig. 3

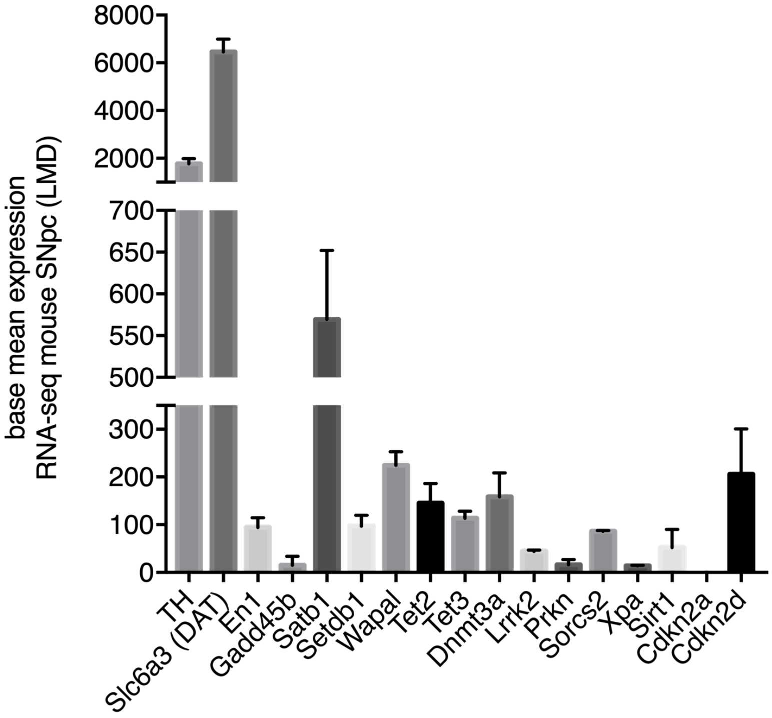
base mean expression
RNA-seq mouse SNpc (LMD)
8000
6000
4000
2000
700
650
600
550
500
300
200
100
0
TH
Slc6a3 (DAT)
En1
Gadd45b
Satb1
Setdb1
Wapal
Tet2
Tet3
Dnmt3a
Lrrk2
Prkn
Sorcs2
Xpa
Sirt1
Cdkn2a
Cdkn2d

Suppl. Fig. 4
